# Intestinal fibroblast heterogeneity: unifying RNA-seq studies and introducing consensus-driven nomenclature

**DOI:** 10.1101/2024.12.09.627573

**Authors:** Neda Glisovic, Aleksandra Chikina, Noémie Robil, Sonia Lameiras, Danijela Matic Vignjevic

## Abstract

Graphical abstract

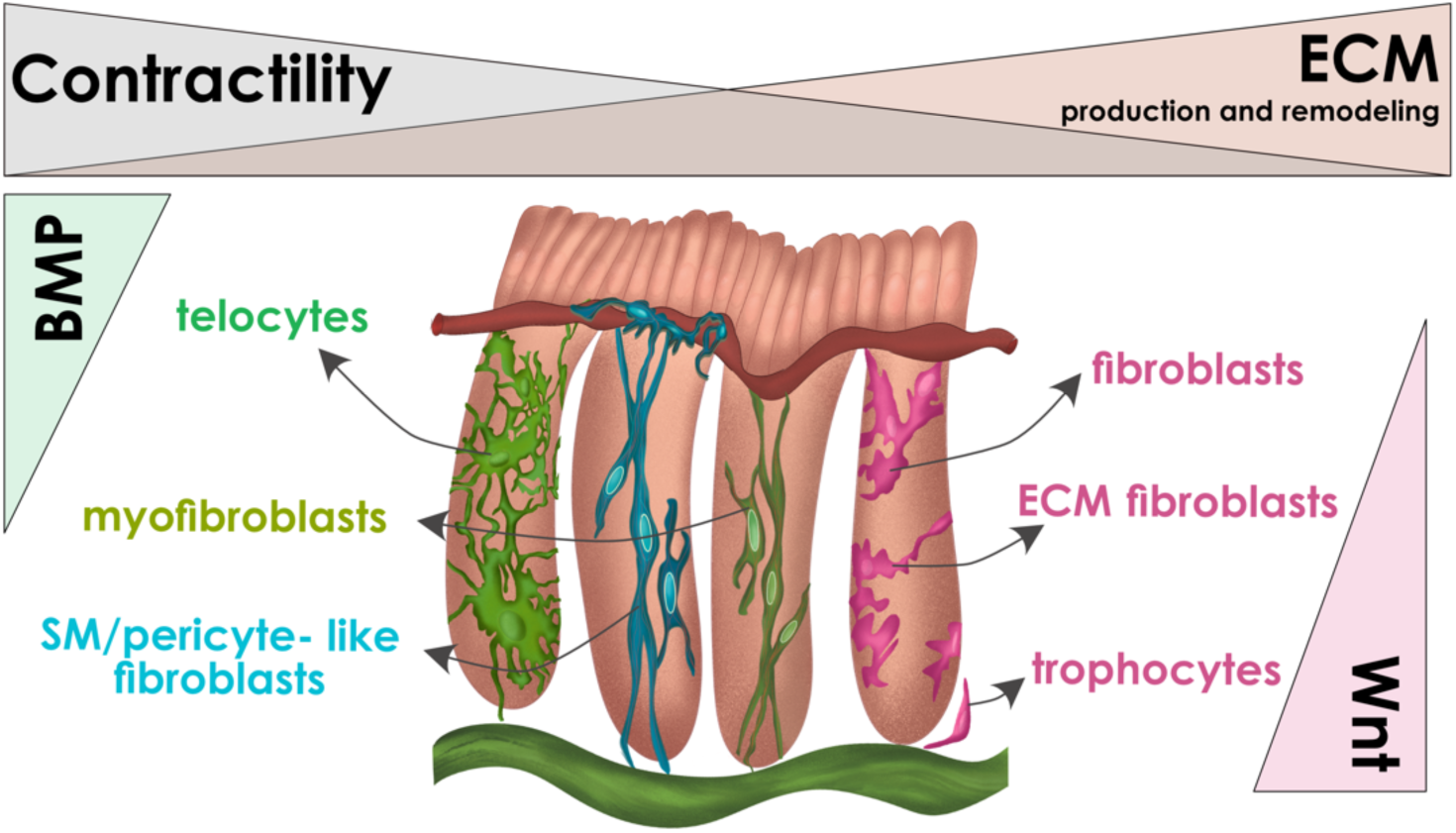

In the colon, single-layered epithelium lines the crypts, which descend into the underlying mucosa. Colonic crypts are surrounded by a network of fibroblasts essential for various functions, including the production and remodeling of the extracellular matrix (ECM), supporting the epithelial stem cell niche, and promoting the differentiation of epithelial cells. Recent studies indicate that fibroblasts are not a homogeneous cell population. However, differences in nomenclature and a lack of exact markers hindered their classification and functional understanding. Here, we used single-cell RNA-sequencing (scRNA-seq) to identify six distinct fibroblast subpopulations in mouse colonic mucosa, each with unique molecular signatures and functional specialization. Our analysis reveals that some fibroblasts are primarily involved in ECM production and remodeling, while others exhibit high contractility. Additionally, a subset of fibroblasts produces cytokines that promote epithelial cell differentiation, whereas another group secrete cytokines essential for maintaining the epithelial stem cell niche. We also map the spatial distribution of these fibroblast subpopulations within the colonic mucosa. Differentiation trajectory analysis suggests distinct pathways for fibroblast differentiation, while cell cycle scoring reveals that fibroblasts do not proliferate under homeostatic conditions. Furthermore, we integrated our scRNA-seq data with previously published datasets to identify common fibroblast populations and propose a standardized nomenclature for intestinal fibroblasts. This unified framework aims to improve communication within the research community and enhance understanding of fibroblast roles in gut homeostasis and gastrointestinal diseases.

## Introduction

The gastrointestinal system consists primarily of the small intestine and colon. Whereas the epithelium aligns villi and crypts in the small intestine, significantly longer crypts in the colon descend into the stroma (*lamina propria*) beneath and reach the first muscle layer (*muscularis mucosae*). At the base of the crypts reside stem cells, perpetually dividing to give rise to transit-amplifying (TA) cells that further differentiate into specialized cell types, including absorptive enterocytes and secretory cells such as goblet, enteroendocrine, and Tuft cells^1^.

Underlying the epithelial cells’ basal side lies the basement membrane, a thin, dense sheet-like structure composed of extracellular matrix (ECM) proteins such as laminin and collagen IV^2^. The basement membrane facilitates epithelial cell adhesion and separates the epithelium from the stroma ^3^. Stroma is found beneath the basement membrane, and it is composed of a network of ECM proteins, such as collagens and fibronectin. It contains mesenchymal cells, including fibroblasts, pericytes, endothelial cells, neurons, glial cells, and immune cells^4,5^. Single-cell RNA sequencing and spatial transcriptomics advancements have unveiled diverse subpopulations of subepithelial fibroblasts^6–9^. These fibroblasts play crucial roles in inflammation, wound repair, and maintaining the epithelial stem cell niche and cell differentiation by secreting ECM, proteases, growth factors, and cytokines, including Wnts, BMPs, and Notch signaling factors^8,10–12^.

Despite considerable research on gut subepithelial fibroblast diversity, there is a need to unify and classify the data. Different studies often assign distinct names to identified fibroblast clusters, even when discussing similar populations^11^. The lack of precise localization, consistent roles, and a single marker for cell population identification complicates the study of subepithelial fibroblasts, resulting in varied nomenclature and confusion. Here, we summarize the characteristics of known intestinal fibroblast populations and propose distinct nomenclature to establish consensus within the field.

## Results

### Distinct subpopulations of fibroblasts in the colonic mucosa

To investigate fibroblast populations in the colonic *lamina propria*, we dissected colons from αSMA:CreER^T2^; R26^mT/mG^ mouse^13,14^, mechanically discarded the muscle layers, sorted live cells, and performed single-cell RNA sequencing. In this mouse model, all cells have red membranes except for cells where the αSMA promoter is active (thus conventional myofibroblasts^15^). Their membranes appear green (EGFP+). To broadly target fibroblast populations, we applied bioinformatics gene-expression filters, selecting cells expressing Vimentin, not expressing H2-Ab1 (MHCII^+^ immune cells) and Krt8 (thus colonic epithelium^16^) and with mitochondrial gene expression ≤20% to exclude potentially damaged cells for more reliable downstream analysis **(Figure 1A).** This approach allowed us to delineate six distinct subpopulations of fibroblasts, denoted as clusters 0, 1, 2, 3, 4, and 5 **(Figure 1B)**. Notably, these encompassed three subpopulations of EGFP+ cells, located in clusters 2, 3, and 4 **(Figure 1B, D).**

**Figure 1.**
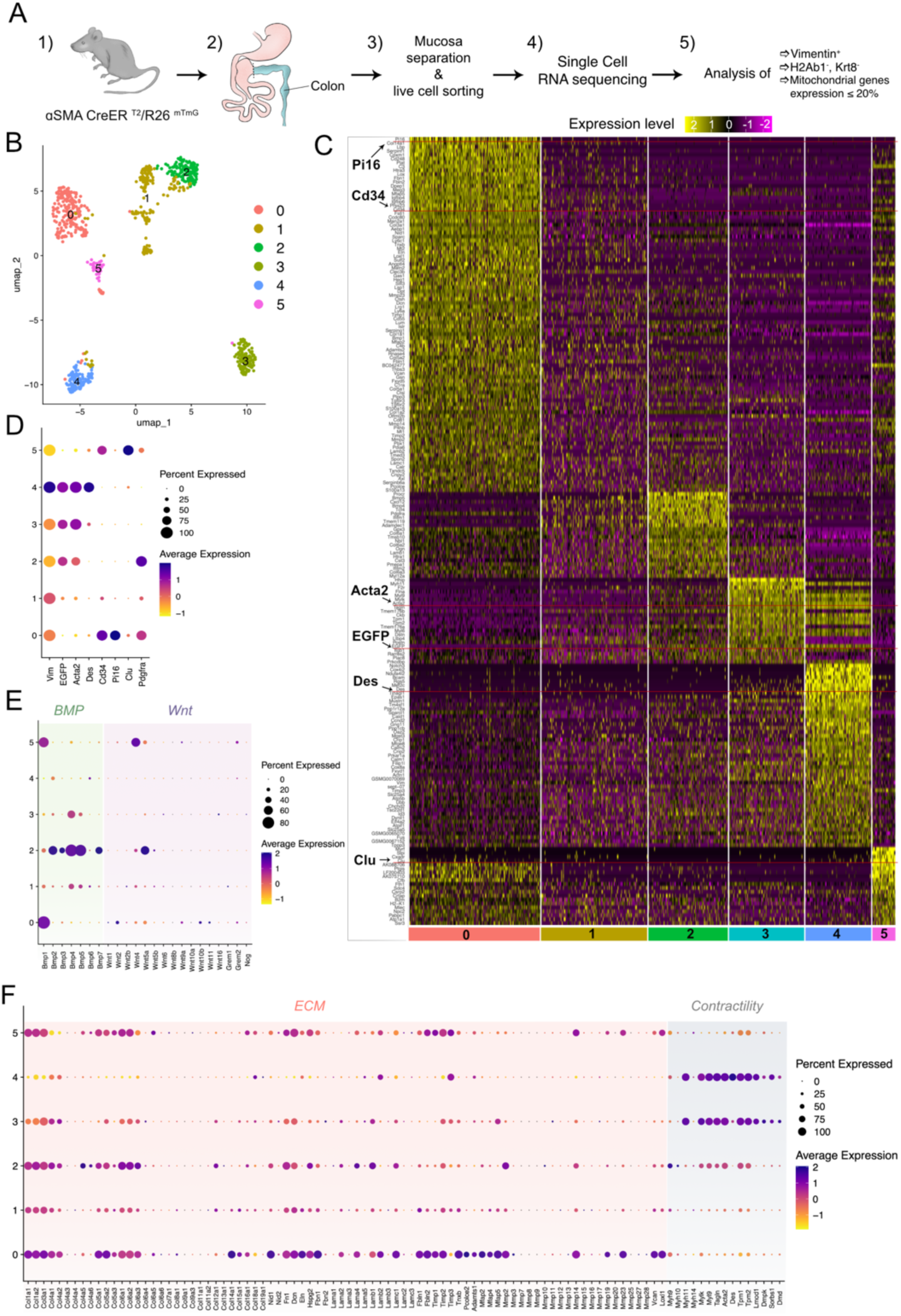
scRNAseq analysis of mouse colonic fibroblasts. **A)** Experimental procedure. Five αSMA:CreER^T^^2^; R26^mT/mG^ mice were used to isolate colons (1,2). Mucosa was physically separated from the muscle layers, live cell sorting was done, and scRNA-Seq was performed (3,4). After, the dataset was enriched in fibroblast-like cells by filtering cells that expressed Vimentin, not expressing H2-Ab1 and Krt8, and with mitochondrial gene expression ≤20% (5). **B)** The UMAP plot of six fibroblast clusters (number of PCs=20). **C)** Heat-map of strongly expressed genes among fibroblast clusters. Marker genes of each of the clusters are highlighted (*Cd34, Acta2, Des, EGFP, Pi16*, and *Clu*). **(D-F)** Expression of selected marker genes among fibroblast clusters. The size of the dot represents the percentage of cells that express the transcript, while the color of the dot is the average expression level within a cluster. **D)** A dot plot of the relative expression of cluster marker genes. **E)** A dot plot of the relative expression of BMP and Wnt pathway genes. **F)** A dot plot of the relative expression of ECM and contractility genes.

To ascertain unique molecular signatures for each fibroblast subpopulation, we calculated the best markers for each population (logFC=1, genes expressed in at least 75% of cells, with p_val_adj <= 0.05), and visualized their expression by generating a heatmap **(Figure 1C)**. This approach allowed us to identify genes prominently expressed in a particular cluster while showing low or negligible expression in others. We delineated distinct molecular signatures for all clusters except cluster 1. In particular, a combination of a few markers was sufficient to distinguish those clusters **(Figure 1D)**. Clusters 0 and 5 were characterized by high expression of *Cd34*, with a notable absence of *Egfp.* They could be further distinguished by the expression of *Pi16* (Cluster 0) or *Clu* (Cluster 5).

In contrast, clusters 2, 3, and 4 expressed *Egfp* and lacked *Cd34*. They could be further distinguished by the expression of *Pdgfra* and low expression of *Acta2* (coding for αSMA *-* alpha-smooth muscle actin) (Cluster 2), high expression of *Acta2* (Cluster 3) and high expression of *Acta2* in addition to *Des* (desmin, an intermediate filament protein) (Cluster 4) **(Figure 1D)**. Cluster 1 presented a challenge as it lacked specific markers that could distinctly differentiate it from the other clusters. It exhibits varied expression levels of all the genes that characterize other clusters, suggesting its transitional nature. Thus, it was excluded from functional analysis.

In the subsequent analysis phase, we used the capabilities of g: Profiler ^17^, an extensive webserver dedicated to functional enrichment analysis. Specifically, we employed g:GOSt within this platform, which compares a custom list of genes provided by the user against various established functional information sources, including Gene Ontology (GO), Kyoto Encyclopedia of Genes and Genomes (KEGG), Reactome, and WikiPathways (WP) (see **Suppl. Table 1** for the analysis links). g:GOSt identifies statistically significant biological processes, pathways, regulatory motifs, and protein complexes. It encompasses three biological concepts: Biological Process (BP), Molecular Function (MF), and Cellular Component (CC) ^18^. For this analysis, we generated gene lists for each cluster, setting the *log.fc* of gene expression to 0.5, and selected only up-regulated genes in each cluster (**Suppl. Table 1).**

Analysis showed that **Cluster 0** (*Egfp*^-^, *Cd34*^+^, *Pi16*^+^) consists of cells enriched for GO terms such as “Extracellular region” (GO:0005576), “Extracellular matrix structural constituent” (GO:0005201), but also “BMP signaling pathway” (GO:0030509). This cluster had a high level of expression of the main constituents of the basement membrane and ECM **(Figure 1F)**. Moreover, a proportion of cells of this cluster expressed high levels of canonical Wnt molecule, *Wnt2*, as well as *Grem2* (**Figure 1E**). This suggests that Cluster 0 is involved in the secretion and maintenance of the basement membrane and ECM, as well as the regulation of the epithelial stem cell niche. **Cluster 5** (*Egfp*, *Cd34*^+^, *Clu*^+^) consists of cells enriched for GO terms such as “Extracellular region” (GO:0005576), “Developmental process” (GO:0032502), “Cell adhesion” (GO:0007155). This cluster was similar to Cluster 0 and, in addition, was characterized by the production of *Wnt4*. Interestingly, clusters 0 and 5 produce *Bmp1*, which mediates the cleavage of pro-collagens and facilitates the assembly of fibrilar collagens ^19–21^ (**Figure 1E**).

**Cluster 2** (*Egfp*^+^, *CD34*^-^, *Acta2^low^*, *Des*^-^) consists of cells enriched for GO terms such as “Extracellular region” (GO:0005576), “Anatomical structure morphogenesis” (Go:0009653), “Extracellular matrix structural constituent” (GO:0005201). Compared to Cluster 1, it contained lower levels of ECM and basement membrane protein transcripts. In addtion to *Col4a1, Col4a2, Lama4, Lama2, Lamb1, Lamc1* that were also expressed in Clusters 0 and 5, *Col4a5, Col4a6, Col12a1* were specifically elevated in Cluster 2 (**Figure 1F**). Cluster 2 also expressed high levels of many BMP transcipts (*Bmp2, Bmp3, Bmp4, Bmp5* and *Bmp7*) as well as non-canonical Wnt, *Wnt5a*. This suggests that Cluster 2 plays a role in epithelial differentiation and ECM production. **Cluster 3** (*Egfp^+^, CD34^-^, Acta2^high^, Des^-^*) was enriched in the following GO terms: “Actin cytoskeleton” (GO:0015629), “Tissue development: (GO:0009888), and “Contractile fiber” (GO:0043292). It expressed high levels of contractility genes, such as myosins and tropomyosins, indicating that Cluster 3 are highly contractile cells. **Cluster 4** (*Egfp^+^, CD34^-^, Acta2^high^, Des^+^*) consists of cells that are highly enriched for GO terms “Muscle structure development” (GO:0061061). The presence of desmin suggests this cluster is more similar to smooth muscle cells and pericytes ^22^ (**Figure 1F**).

Overall, this analysis indicates that, consistent with previous observations, subepithelial fibroblasts express genes involved in ECM remodeling and cytokines that regulate the fate of epithelial cells, including stemness, proliferation, and differentiation ^10–12,23^. However, these functions were distributed across different clusters, suggesting a specialization of fibroblasts for specific roles. For instance, Clusters 0 and 5 primarily focused on ECM production, while Clusters 3 and 4 showed high expression of genes related to contraction (**Figure 1F**). This specialization is noteworthy because fibroblasts typically perform both functions simultaneously ^12^. Additionally, there was a functional division in the production of BMP/Wnt molecules. Cluster 2 mainly contributed to the production of BMPs, indicating a role in epithelial differentiation (**Figure 1E**). In contrast, Clusters 0 and 5 appeared to support the maintenance of the epithelial stem cell niche by producing Wnt molecules and BMP antagonists.

### Spatial organization of fibroblast subpopulations in the colon mucosa

Despite the wealth of studies on the diversity and heterogeneity of gut fibroblasts, a comprehensive understanding of the spatial distribution and visualization of various fibroblast types remains elusive. Questions about where different populations localize and whether this localization correlates with their functional roles remain to be further investigated ^11^. To address this, we performed multicolor imaging, which enabled us to visualize and spatially map different fibroblast subpopulations identified in our RNA sequencing analysis.

The initial step in distinguishing between clusters involved identifying cells as either EGFP^+^ or CD34^+^ (**Figure 2A, B**). In EGFP^+^ cells, we assessed the expression of αSMA protein derived from the *Acta2* gene. Cells exhibiting low αSMA protein levels were categorized into Cluster 2 (EGFP^+^, αSMA^low^). In the mouse line we used, tamoxifen-dependent Cre activity is driven by a 1.1 kb fragment of the mouse *Acta2* promoter ^14^. While this promoter fragment is sufficient for CreER^T2^ expression and subsequent excision of a loxP-flanked *Tomato* reporter cassette (rendering the cells EGFP+), the endogenous transcriptional machinery in Cluster 2 cells is not robust enough to fully activate the native *Acta2* gene. Consequently, these cells exhibit low levels of αSMA protein. Desmin expression was then used to further delineate cells with high levels of αSMA protein. Those with low desmin levels were allocated to Cluster 3 (EGFP^+^, αSMA^high^, desmin^low^), while cells with high desmin levels were placed in Cluster 4 (EGFP^+^, αSMA^high^, desmin^high^). Clusters 0 and 5 were characterized by CD34 expression without EGFP. Despite identifying *Pi16* and *Clu* transcripts as specific markers for Clusters 0 and 5, respectively (**Figure 1C, D**), commercially available antibodies did not yield satisfactory results, preventing a clear distinction between Clusters 0 and 5.

**Figure 2.**
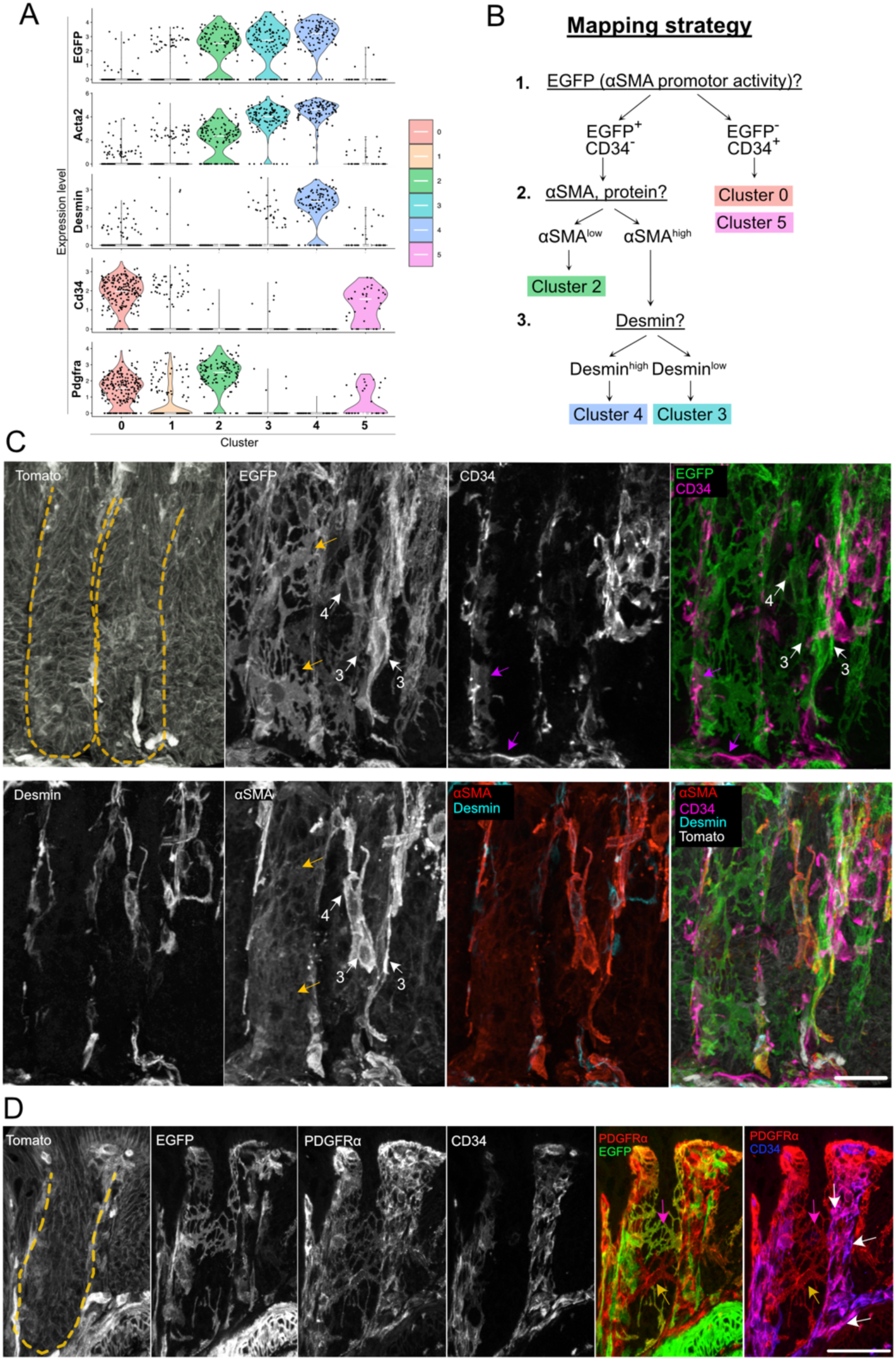
Spatial distribution of colonic fibroblasts. **A)** Violin plots of the expression of specific fibroblast markers across different fibroblast clusters. The x-axis identifies the fibroblast clusters, and the y-axis shows the level of gene expression for each marker. **B)** Mapping strategy used for spatial visualization of different fibroblast clusters. **(C and D)** Immunofluorescence of the distal colon sections from αSMA:CreER^T^^2^; R26^mT/mG^ mice (scale bar, 40 μm). Cell membranes of αSMA+ fibroblasts are labeled with EGFP, all the other cell membranes are labeled with Tomato. Crypts are outlined with yellow dashed lines. **C)** Tomato, EGFP, staining of desmin, αSMA protein, CD34. Yellow arrows mark stellate cells of Cluster 2 (EGFP^+^, CD34^-^, desmin^-^, αSMA^low^). White arrows point to cells of Clusters 3 and 4 (annotated with the number that matches the cluster number). While all three cells showed high expression of EGFP and αSMA protein, the cell annotated with number 4 has strongly stained desmin fibers, indicating it belonged to Cluster 4. Cells annotated with number 3 showed weak staining of desmin, suggesting they were part of Cluster 3. Purple arrows point to CD34^+^ cells (Clusters 0 and 5). **D)** Tomato, EGFP, PDGFRα, CD34 staining. The purple arrow indicates EGFP^+^, PDGFRα^+^ (Cluster 2). Yellow arrows indicate PDGFRα^+^(cluster 1). White arrows show CD34^+^, PDGFRα^+^ cells (Clusters 0 and 5)

Using this mapping strategy, we successfully visualized distinct clusters of subepithelial fibroblasts in the distal (**Figure 2C**) and proximal sections of the murine colon (**Suppl. Figure 1A-C**). Cluster 2 was characterized by EGFP^+^ stellate-shaped (branched) cells, interconnected via long, thin dendritic protrusions (**Figure 2C**, **Figure 3A-C, Suppl. Movies 1-3**) and gap junctions (connexin 43, gene *Gja1*) to form a network encapsulating the crypt (**Suppl. Figure 2E-F**). Their morphology was consistent across various gut segments. Clusters 0 and 5 similarly exhibited a stellate-like morphology but with fewer, thicker protrusions that taper to relatively thin ends, connecting to adjacent cells by connexin 43 (**Figure 2C, Suppl. Figure 2E-F, Suppl. Movie 2-3**). These clusters were found along the entire length of the crypts, often in tightly packed groups, making individual cell identification challenging (**Figure 2C**).

**Figure 3.**
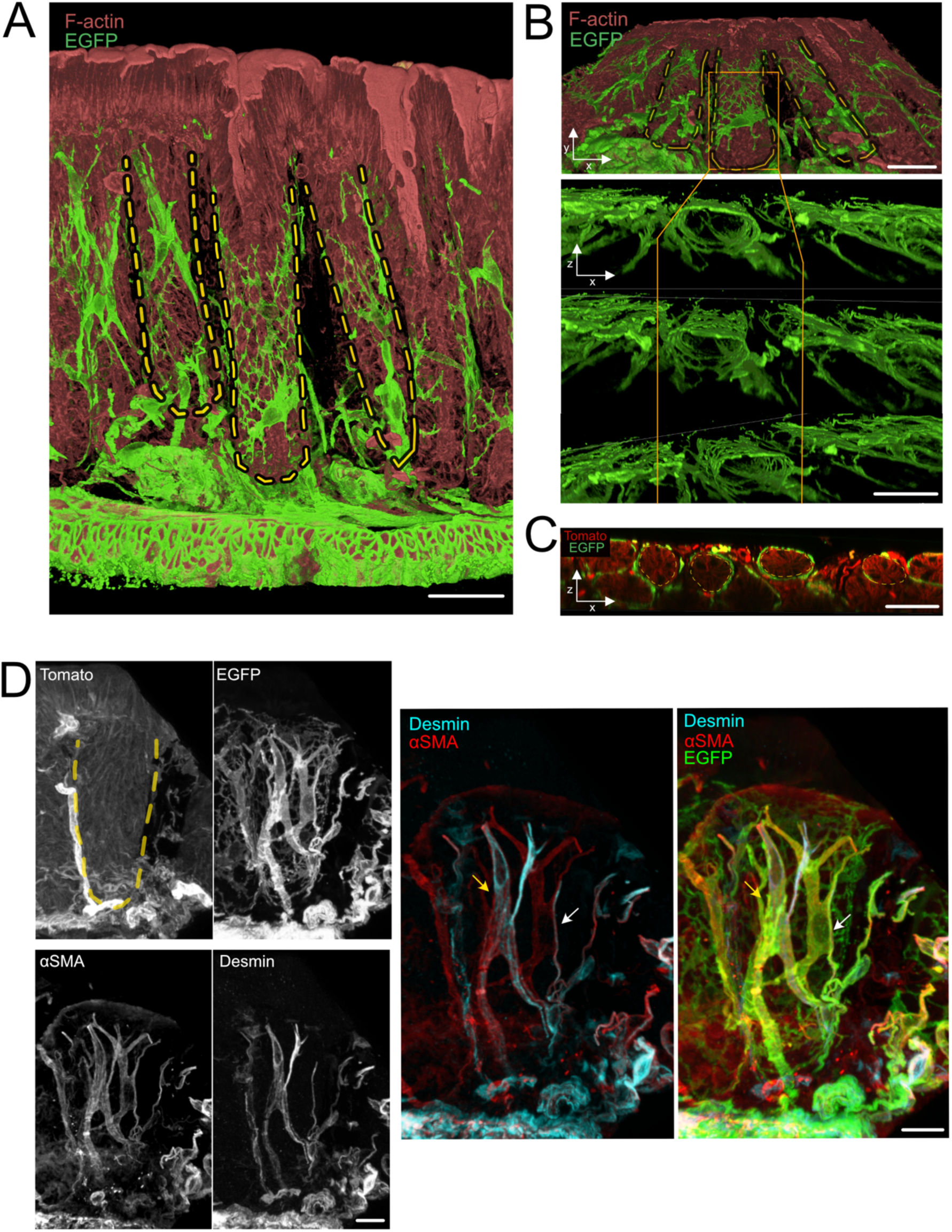
Spatial distribution of EGFP+ fibroblasts. Immunofluorescence of the distal colon from αSMA:CreER^T^^2^; R26^mT/mG^ mice (scale bar, A-C: 40 μm, D: 20 μm). Cell membranes of αSMA+ fibroblasts were labeled with EGFP, all the other cell membranes were labeled with Tomato. Crypts are outlined with yellow dashed lines. **A)** Three-dimensional representation of longitudinally sectioned crypts. F-actin is depicted in red, and αSMA^+^ fibroblasts are shown in green (EGFP). **B)** Upper panel: a top-down view of the crypts shown in A (x-y plane), lower panel: orthogonal z-sections of the same crypts (x-z plane). Red (F-actin), and αSMA^+^ fibroblasts in green (EGFP). **C)** Orthogonal Z-sections (x-z plane) of the crypts depicted in A. αSMA^+^ fibroblasts were visualized in green (EGFP), while all other cell membranes are in red (Tomato). **D)** Tomato, EGFP, staining of desmin and αSMA protein. White arrow points to EGFP^+^ αSMA^high^ Desmin^low^ cell (Cluster 3). Yellow arrow highlights the EGFP^+^ αSMA^high^ Desmin^high^ cell (Cluster 4).

Despite CD34 being a marker for endothelial progenitor cells, we distinguished fibroblasts from blood vessels by their unique location, morphology, and the absence of CD31 staining, which is specific to blood vessels ^24^ (**Suppl. Figure 2A-B, Suppl. Figure 1B**). Clusters 3 and 4 were primarily composed of spindle-shaped cells, varying from smaller, square-like cells similar in size (**Figure 2C**, **Figure 3A, Suppl. Movie 1**) to exceptionally long cells connecting the *muscularis mucosae* at the crypt base to blood vessels at the top (up to 120 μm long, **Figure 3D, Suppl. figure 3A, B**). These cells, found on top or in between crypts, were either solitary or in small groups of 3-4 cells. Both clusters featured cells with prominent αSMA-rich contractile stress fibers, with cluster 4 also showing an abundance of desmin fibers (**Figure 2C**, **Figure 3D, Suppl. Figure 3A, B**). When found together, EGFP^+^ fibroblasts (Clusters 2-4) were positioned directly adjacent to the basement membrane of epithelial cells, labeled with as pan-laminin (LAM1), COL1, COL4, and FN1 (**Suppl. Figure 4A-E**). In contrast, Clusters 0 and 5 were situated further from the epithelial cells, residing above the EGFP^+^ fibroblasts. However, when alone, cells from clusters 0 and 5 interacted directly with the epithelial basement membrane (**Suppl. Figure 2G**).

We also examined PDGFRα expression, a frequently cited fibroblast marker. PDGFRα expression was most significant in Cluster 2 (EGFP^+^), followed by Cluster 0 (CD34^+^), and absent in Clusters 3 and 4 (**Figure 1D, 2A**). Consistent with literature reports, we observed a PDGFRα expression gradient along crypts ^8^ with cells atop the crypts, including pericytes over blood vessels, exhibiting higher PDGFRα levels than those at the bottom (**Figure 2D, Suppl. Figure 1C**). Intriguingly, we also identified cells negative for CD34 and EGFP but positive for PDGFRα, suggesting a potential association with Cluster 1 based on their morphology and positioning similar to Cluster 2 (**Figure 2D, Suppl. Figure 1C**). The UMAP plot shows that a small proportion of cells of Cluster 1 express PDGFRα (**Supp.** Figure 1D). The expression of intermediate filament vimentin was observed in all fibroblast clusters (**Figure 1D, Suppl. Figure 2C-D**), but it followed the same pattern as PDGFRα, with higher expression at the top of the crypt and lower expression in fibroblasts closer to the crypt bottom (**Suppl. Figure 2C**).

Altogether, we found that fibroblasts exhibit distinct shapes and localizations in the colonic mucosa. Stellate-shaped fibroblasts were located close to the epithelium, forming a basket around the crypts. These are enriched in ECM genes and produce cytokines necessary for the maintenance of the stem cell niche (Clusters 0 and 5) or cytokines that support the differentiation of epithelial cells (Cluster 2). In contrast, spindle-shaped cells do not express ECM genes, but are highly enriched in genes related to contractility (Cluster 3, Cluster 4). The spindle cells can be attached on one side to blood vessels at the top of the crypts just beneath the epithelium and on the other side to the muscle layer. In addition to expressing contractility genes, some of the spindle cells are enriched in genes typically found in pericytes and smooth muscle cells (Cluster 4).

### Different fibroblast subpopulations were at various stages of differentiation

To investigate fibroblast differentiation pathways based on gene expression profiles, we employed the Monocle 3 algorithm. Cluster 1 was selected as the root for trajectory analysis due to its expression of genes present in all other clusters (**Figure 1C**). The obtained pseudotime trajectory was projected onto UMAPs, and cells were colored according to Seurat fibroblast clusters and predicted pseudotime values, respectively (**Figure 4A, B**). Each leaf, represented by light gray circles, indicates a unique outcome (cell fate or end state) of the trajectory. Black circles denote main branch nodes, which direct cells toward one of several potential fates. Eight distinct branches were identified (**Figure 4A, B**).

**Figure 4.**
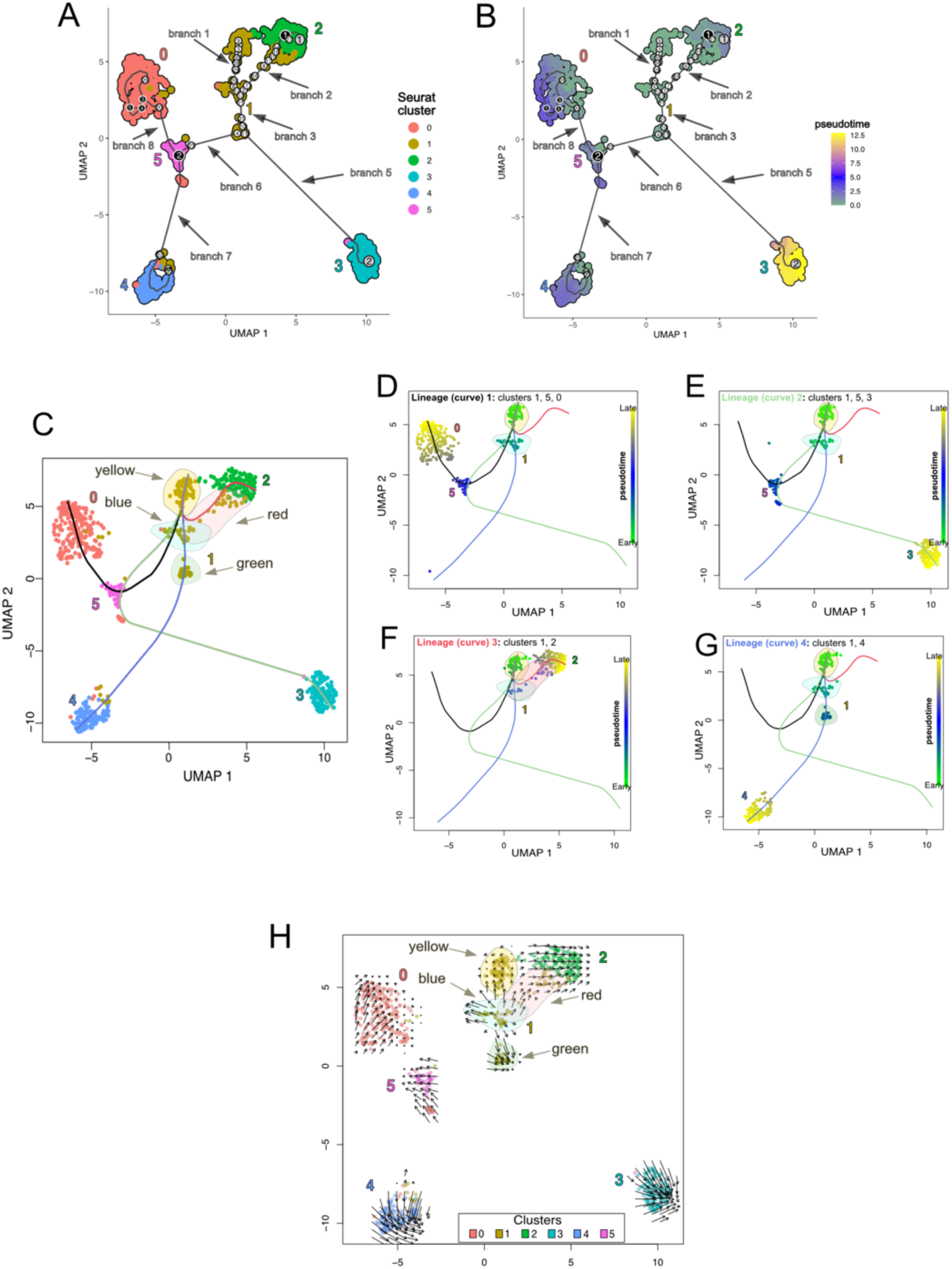
Pseudotime trajectory analysis of murine colonic fibroblasts. (**A** and **B**) Pseudotime trajectory, projected on UMAPs, modeled by Monocle 3. Cells are coloured according to Seurat fibroblast clusters and predicted pseudotime, respectively. The circles with numbers inside signify different spots on the trajectory. Each leaf, represented by light grey circles, represents a unique outcome (cell fate) of the trajectory. The black circles represent branch nodes, which allow cells to proceed to one of several fates. The trajectory is divided into branches numbered 1 through 8. **C**) Lineage trajectories (black, green, red, and blue lines) predicted by Slingshot, projected onto UMAP. Cells are colored according to Seurat fibroblast clusters. Four clouds of cluster 1 cells are labeled yellow, red, blue, and green, and highlighted in the same colors. **D-G**) Lineage trajectories (black, green, red, and blue lines/curves) predicted by Slingshot, projected onto UMAPs. Cells are colored according to the predicted pseudotime. **H**) UMAP representation of 6 fibroblast clusters with RNA velocity vectors. Fibroblast clusters are color-coded by Seurat fibroblast clusters. Four clouds of cluster 1 cells are labeled yellow, red, blue, and green, and highlighted in the same colors.

In Cluster 1, three branches contained cells with low pseudotime values (**Figure 4A, B**). Branch 2 terminated at a leaf (gray circle 1) corresponding to Cluster 2. The pseudotime values of cells along Branch 3 increased, suggesting differentiation toward Cluster 2. Branch 3 subsequently divided into Branch 5, which terminated at a leaf (gray circle 2) corresponding to Cluster 3. Cells in Cluster 3 exhibited the highest pseudotime values, indicating their status as the most differentiated and well-defined population.

Branch 6 divided at the major branching point 2 (black circle) into two pathways: Branch 7, terminating in Cluster 4, and Branch 8, concluding in Cluster 0. Pseudotime values along these branches increased progressively, reflecting differentiation toward the well-defined clusters 0 and 4. Interestingly, the major branching point 2 was located in Cluster 5, suggesting that cells in Cluster 5 can transition toward Cluster 4 or Cluster 0. Nevertheless, cells in Cluster 5 displayed a gradient of pseudotime values (**Figure 4A, B**).

To complement this, we performed diffusion pseudotime analysis using the R package Slingshot, designating Cluster 1 as the root (**Figure 4C-G**). Four distinct differentiation trajectories were identified and projected onto UMAPs, leading from Cluster 1 to Cluster 0 or 3 via Cluster 5, or directly to Clusters 2 and 4 (**Figure 4C-G**). Cells were colored according to Seurat fibroblast clusters (**Figure 4C**) or predicted pseudotime values (**Figure 4D-G**). Within Cluster 1, four subpopulations (yellow, red, blue, and green) were detected (**Figure 4C-G**). Cells in the yellow and blue subpopulations originated all trajectories, while red subpopulation cells followed the trajectory leading to Cluster 2 (corresponding to Branch 2 in Monocle 3; **Figure 4A, B**). Green subpopulation cells were restricted to the trajectory leading to Cluster 4 (Slingshot; **Figure 4G**) and were located at the convergence of Branches 3, 5, and 6 (Monocle 3; **Figure 4A, B**).

We also employed RNA velocity analysis using the R package velocyto. RNA velocity analysis predicts the future state of single cells based on ratios of spliced, unspliced, and degraded mRNA, visualized as velocity vectors projected onto UMAPs (**Figure 4H**). Cells were colored by Seurat fibroblast clusters. Short arrows corresponded to terminally differentiated cells, while long arrows reflected significant gene expression changes. Cells differentiated along the direction of these arrows. Cluster 1 exhibited a distinct organization compared to other clusters (**Figure 4H**). It appeared as a central hub, with cells displaying opposing long and short RNA velocity vectors, potentially indicating active differentiation in multiple directions. Examining the four subpopulations within Cluster 1 (yellow, red, blue, and green) revealed that red subpopulation cells pointed toward Cluster 2, consistent with both trajectory analyses. The velocity vectors of yellow and blue subpopulation cells diverged, indicating differentiation toward distinct end states. Green subpopulation cells’ velocity vectors pointed toward Cluster 3. Additionally, velocity vector lengths varied within Clusters 2 and 5, aligning with their end states and increasing pseudotime values (**Figure 4H, B**). Clusters 3 and 4 showed long, unidirectional velocity vectors, indicating active differentiation toward their final states, again corresponding to higher pseudotime values (**Figure 4H, B**).

Trajectory analyses and RNA velocity data collectively suggest that Cluster 1 is composed of multipotent cells. The central location, low pseudotime values, and opposing RNA velocity vectors support its role as a reservoir of precursor cells for different fibroblast clusters.

### Subepithelial fibroblasts do not proliferate

To determine whether fibroblasts proliferate, we used the Cell Cycle Scoring function in R (Seurat, version 4.3) to calculate potential cell cycle scores for each cluster based on the expression of classical marker genes for the G2/M and S phases ^25^. We observed that the S and G2-M scores were close to zero across all clusters, except Cluster 5, where a subset of cells displayed slightly higher score levels (**Suppl. Figure 5A,B**). Furthermore, genes linked with various cell cycle phases showed no or extremely low expression across the clusters except Cluster 5, and slightly less in Cluster 0. (**Suppl. Figure 5C**). Altogether, this suggests that fibroblasts in homeostasis do not proliferate excessively. Ki67 immunostaining furhter confirmed that fibroblasts do not divide (**Suppl. Figure 5D**). This agrees with Kinchen et al., who show that most fibroblasts are in the G0/G1 phase ^7^.

### Integration of data with already published datasets

Next, we examined how the fibroblast populations we have identified relate to previously published datasets (Degirmenci et al., Kinchen et al., McCarthy et al., and Roulis et al.) ^6–9^. To compare cluster definitions, each dataset has been reanalyzed using the publication thresholds and parameters. Finally, by comparing markers, we redefined clusters from the original analysis. After suppressing immune clusters from Kinchen and McCarthy datasets using *Ptprc* (CD45 protein) expression as a marker, all datasets were integrated with the Seurat 3.2 CCA algorithm. This integrated dataset was clustered in the same way as described above, using 15 PCs and a resolution parameter of 0.8, defining 19 clusters (**Figure 5A**).

**Figure 5.**
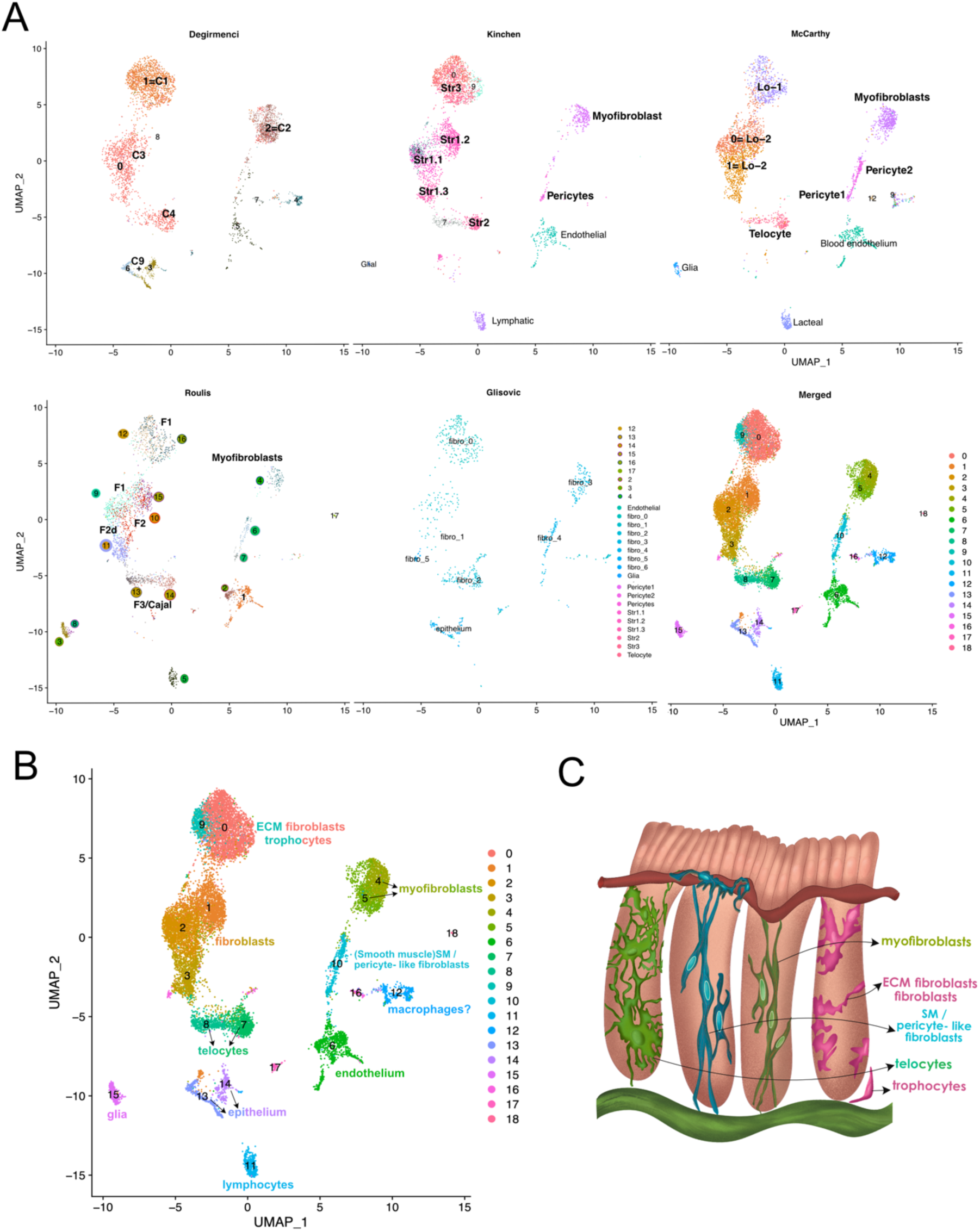
Integrated scRNAseq analysis of intestinal fibroblasts. (A) UMAP representation of different studies, integrated using CCA algorithm from Seurat and split by individual datasets (Degirmenci et al., Kinchen et al., McCarthy et al., Roulis et al., and Glisovic et al.). The last UMAP represents an integrated UMAP plot, with all the cells from all the studies. Clustering analysis resulted in 19 clusters. (B) Integrated UMAP plot with the assigned names to distinct clusters. (C) Fibroblasts in the mouse colon are categorized into fibroblasts and ECM fibroblasts/ trophocytes (in pink), then myofibroblasts, smooth muscle/ pericytes-like fibroblasts, and telocytes (in different shades of green).

Individual data sets were then overlayed on the unified map to see how subpopulations of fibroblasts identified in individual studies relate to new clusters (**Figure 5A, B**). Based on molecular signatures from 19 identified clusters, we can exclude 6 clusters as they are identified as epithelial, endothelial and glial cells, macrophages and lymphocytes. Ten other clusters were identified as fibroblasts according to gene signatures reported across studies. Three small clusters (16, 17 and 18) could not be matched to any already described cell type. Interestingly, cluster 17 was enriched in cell cycle genes, suggesting it contains cells that divide (**Suppl. Figure 5E**).

Several of the remaining 13 clusters have a molecular signature consistent with most studies, such as myofibroblasts (clusters 4 and 5) and telocytes (clusters 6 and 7). Cluster 10 is often annotated as pericytes. However, based on the spatial analysis we have performed here, cells of this cluster were not always associated with the blood vessels. Clusters 0 and 9 represent fibroblasts that are specialized in ECM production and, in some studies, are annotated as crypt bottom fibroblasts involved in stem cell niche maintenance. Clusters 1, 2 and 3 were given different names in different studies. Comparing clusters from individual datasets allowed us to refine the nomenclature.

## Discussion

### Characterization of distinct fibroblast subtypes

Based on gene expression, spatial localization of different clusters, and integration with previous studies, we propose a uniform intestinal fibroblasts classification that includes five subpopulations: telocytes, trophocytes and ECM fibroblasts, fibroblasts, myofibroblasts and smooth muscle/pericytes-like fibroblasts (**Figure 5C**). **Table 1** summarises the marker genes and descriptions for all populations, while their signatures are plotted on the UMAPs in **Suppl. Figure 6**.

**Table 1.**
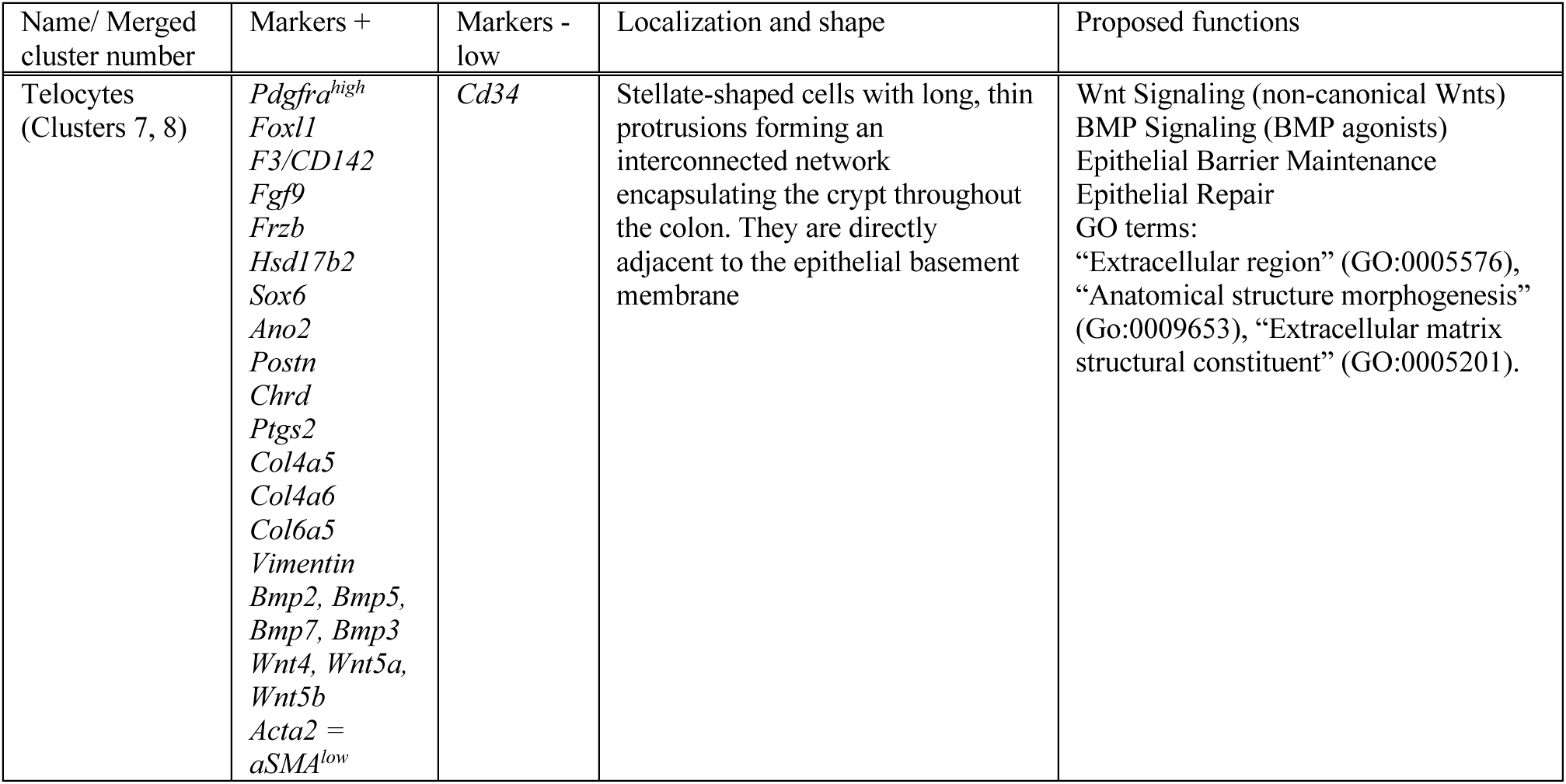
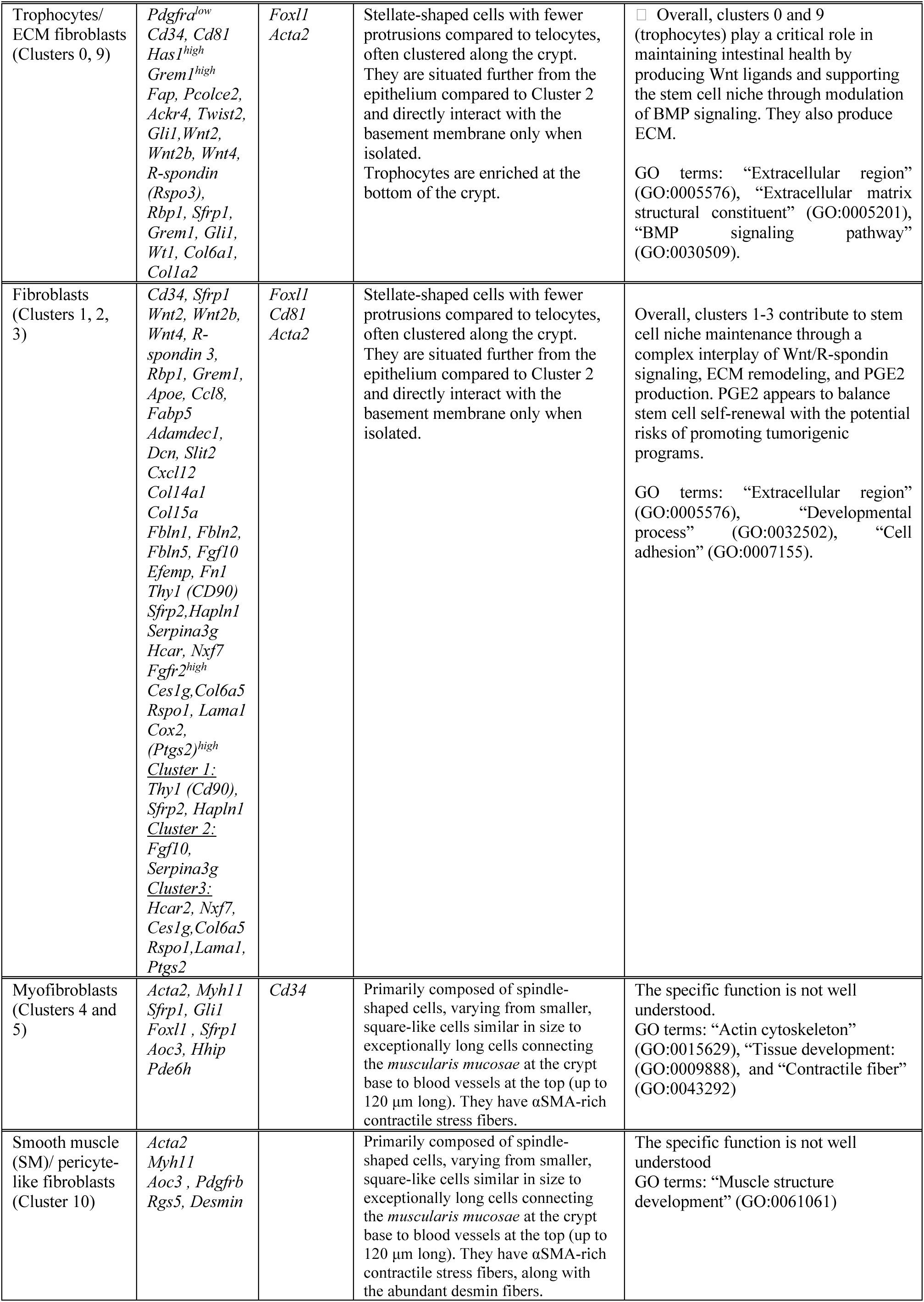
Summary of characteristics of different fibroblast populations.

### Telocytes – clusters 7 and 8

According to the authors of all the studies, telocytes are distinguished by their high *Pdgfrα* expression. Degirmenci et al. (C4) and Kinchen et al. (Str2) also report high *Foxl1* expression, and McCarty et al. points out that they do not express *Cd34* (**Suppl. Figure 7**). While Roulis et al. (F3/Cajal cluster) characterizes telocytes with high *Ano2* expression, Kinchen et al. identify additional markers, including *F3/Cd142*, *Fgf9, Frzb, Hsd17b2,* and the transcription factor *Sox6*. Telocytes are represented in our analysis as cluster 2, and we also found that this cluster has low *Acta2* expression. Both Degirmenci et al. and McCarty et al., as well as this study, identified telocytes as a source of non-canonical Wnt ligands (*Wnt4, Wnt5a*, and *Wnt5b*). However, there’s further complexity in Wnt signaling within the colon. *Pdgfrα*^low^ cells (described later), express the canonical Wnt ligand *Wnt2b* alongside several factors that enhance Wnt signaling (Rspo factors). In contrast, telocytes express primarily non-canonical Wnt molecules (*Wnt4, Wnt5a,* and *Wnt5b*) with lower levels of Rspo factors compared to *Pdgfrα*^low^ cells. Interestingly, both cell types produce Wnt inhibitors (for example, *Frzb* by telocytes), suggesting a tightly regulated interplay in Wnt signaling within the colon.

*Wnt5a*, specifically expressed by telocytes, plays a crucial role in epithelial repair after injury by enhancing TGFβ signaling ^26^. Additionally, Kinchen et al. also observed high periostin (*Postn*) expression in telocytes. While periostin is essential for tissue repair, it has also been linked to tumorigenesis ^27^. This dual functionality of periostin expressed by telocytes highlights the potential for both beneficial and detrimental effects depending on the context.

In contrast to Wnt signaling, transcripts for BMP signaling components exhibit a distinct pattern. Agonists for BMP signaling are predominantly expressed in telocytes (TGFβ superfamily ligands: *Bmp2* and *Bmp5*, then *Bmp7, Bmp3*). Notably, *Bmp7*, a crucial factor for forming potent BMP signaling dimers, is specifically found in telocytes (McCarthy et al). This suggests that telocytes may be a reservoir for colon BMP signaling. Among BMP inhibitors, *Grem1* is restricted to *Pdgfrα*^low^ stromal cells, while *Chrd* is expressed higher in telocytes.

Interestingly, Kinchen et al. point out the existence of 2 subpopulations of telocytes: the first, enriched in GO terms such as “ BMP signaling and response”, and the second, which expresses factors related to “response to wound healing” and “regulation of epithelial cell proliferation”. Roulis’ pathway analysis revealed an increase in arachidonic acid metabolism in telocytes (together with the F2d subcluster, which corresponds to cluster 3, as discussed later). Cyclooxygenase-2 (COX2, also known as *Ptgs2*) is highly expressed in telocytes, strongly supporting their participation in wound healing, given the importance of COX2 in this process. Moreover, Kinchen et al. and Roulis et al, but also our analysis, report the expression of collagens by telocytes (Kinchen: *Col4a5*, *Cola4a6*, Roulis: *Col6a5*), as well as their proximity to the epithelial monolayer, pointing out their role in epithelial barrier maintenance. In the small intestine, telocytes gather at the base of the villus and serve as a BMP reservoir, while *Pdgfrα*^low^ *Grem1*^+^ cells are seen on the bottom of the crypts. This spatial distribution does not appear in the colon. Telocytes are spread throughout the whole crypt length, forming the basket around them. They are similar to cluster 2 in our dataset.

### Trophocytes/ ECM fibroblasts - Clusters 0 and 9

Clusters 0 and 9 of the integrated dataset overlap with Degirmenci’s cluster 1, McCarty’s cluster Lo-1, Roulis’ cluster F1, Kinchen’s cluster Str 3, and our cluster 0. McCarty et al. (Lo-1) classify these fibroblasts as *Pdgfrα*^low^, *Foxl1*^-^, *Cd34*^+^, and *Cd81*^+^, while Roulis notes their strong *Has1* expression (**Suppl. Figure 8, 9**). Kinchen et al. includes a few additional markers in their description: *Fap, Pcolce2, Ackr4*, and *Twist2*, whereas Degirmenci et al characterize them by *Gli1* expression, even though it is widely expressed in all their clusters.

Fibroblasts of clusters 0 and 9 majorly produce canonical Wnt molecules: *Wnt2, Wnt2b, Wnt4*, and R-spondin (*Rspo3*), *Rbp1*, and *Sfrp1*. They highly express *Grem1* (McCarthy et al). Notably, these markers are also present in clusters 1-3, albeit at lower levels and in fewer cells, except *Sfrp1*, which is highly expressed in clusters 1-3.

Several experimental studies have demonstrated the role of these cells in intestinal health and the maintenance of the stem cell niche. McCarty et al. showed that *Grem1*^+^ cells are necessary for intestinal health since their depletion caused severe intestinal damage and lower lifespan in mice. These cells are not required for the Wnt-dependent proliferation of transient amplifying cells, but they are critical for sustaining intestinal stem cells by blocking BMP signaling. *In vitro*, *Pdgfrα*^low^ cells can replace recombinant *Nog* (Noggin) and *Rspo1* to promote crypt development into enteric spheroids ^8^. *Pdgfrα*^low^ *Cd81*^+^ Lo1 cells (McCarthy et al.) were particularly successful at providing the trophic factors required to sustain intestinal stem cells and counter BMP signals.

Furthermore, Degirmenci et al. showed the proximity of *Gli1*^+^*Wnt4*^+^ cells to the base of the crypts, as well as the spatial overlapping of those cells with *Rbp1* and *Sfrp1*. They show that *Gli1*^+^ cells serve as the essential niche for epithelial stem cells of the colon and as a reserve niche for the stem cells in the small intestine.

Interestingly, Kinchen’s trajectory study identified *Wt1* as a marker that segregates inside the Str3 cluster, a likely progenitor population. Kinchen’s GO enrichment analysis also revealed enrichment in ‘‘supramolecular fiber organization’’ and ‘‘extracellular cluster organization’ pathways in this cluster, with significant expression of *Col6a1*, *Col1a2*, and *Pcolce2*, which goes with our observations. They are similar to cluster 0 in our dataset.

### Fibroblasts – Clusters 1, 2, 3

Similar to clusters 0 and 9, clusters 1-3 exhibit several intriguingly distinct characteristics. McCarty (Lo-2) defines cluster 1-3 fibroblasts as *Pdgfrα*^low^, *Foxl1*^-^, *Cd34*^+^, *Sfrp1*^+^ and *Cd81*^-^, which particularly distinguishes them from Clusters 1 and 9 (McCarthy Lo-1, *Cd81*^+^) (**Suppl. Figure 10**). As mentioned before, fibroblasts of clusters 0 and 9 are the major producers of *Wnt2, Wnt2b,* and *Wnt4*, as well as R-spondin (*Rspo3*), *Rbp1, Grem1*, and *Sfrp1* (**Suppl. Figure 9**). Importantly, these markers are present in clusters 1-3 but at lower levels and in fewer cells, except *Sfrp1*, which is more abundant in clusters 1-3.

Clusters 1-3 of fibroblasts are comparable to Kinchen’s cluster Str1. Their GO enrichment analysis revealed enrichment in GO terms such as “positive regulation of locomotion”, “response to tumor necrosis factor”, “ERK1 and ERK2” cascade, and “extracellular matrix” in these cells. *Apoe, Ccl8, Fabp5, Adamdec1, Dcn, Slit2*, and *Cxcl12* were some of the most notably stimulated genes. They also discovered that clusters 1-3 fibroblasts are enriched in non-fibrilar collagens (*Col14a1, Col15a*) and elastic fibers (*Fbln1, Fblin2, Fbln5, Efemp1, Fn1*), unlike telocytes which express sheet collagens (*Col4a5, Col4a6*). Furthermore, Kinchen’s Str 1.2 corresponds to cluster 1 and has intermediate *Thy1* expression (*Cd90*). It also produces *Sfrp2* and *Hapln1*. Kinchen’s Str1.1, which corresponds to cluster 2, expresses *Fgf10* and *Serpina3g*. Kinchen’s Str1.3, which corresponds to cluster 3, expresses *Hcar2* and *Nxf7*.

Furthermore, clusters 1-3 of the integrated dataset are likely to overlap with Roulis cluster F2. They exhibit high expression of *Fgfr2*. Expression of *Ces1g, Col6a5, Rspo1,* and *Lama1* describe a subcluster of F2 called F2d. It is most likely equivalent to the final integrated cluster 3. Pathway study demonstrated increased arachidonic acid metabolism in the F2d subcluster (but also in Roulis’ cluster F3: telocytes). Cyclooxygenase-2 (COX2, also known as *Ptgs2*), responsible for the production of PGE2, is highly expressed in these two clusters. Roulis defines the F2d subcluster as cells near the stem-cell zone. They are referred to as rare pericryptal *Ptgs2*-expressing fibroblasts (RPPFs). Their localization is associated with the expression of *Lama1*, a basement membrane protein, and R-spondin (*Rspo1*), a stem cell niche factor found at the crypt base.

*Col6a5* effectively demarcated the entire 1-3 population. Using *Col6Cre* Ptgs2 ^f/f^ mice, Roulis et al. selectively depleted COX2 in all fibroblasts within this population. Subsequent analysis revealed a significant reduction in *Ptgs2* expression levels throughout the tissue, confirming that fibroblasts 1-3 are the primary source COX2 in intestinal homeostasis. Additionally, they investigated the impact of PGE2 on intestinal stem cell function. Crypts from the small intestine were cultured in organoid growth media supplemented with PGE2. PGE2 inhibited the formation of budding organoids, resulting in spheroid-like structures lacking crypt-villus architecture, indicative of poor differentiation and increased stemness. These spheroid-like structures harbored a higher proportion of stem cells with complete organoid-forming capacity. They further showed that fibroblast-derived PGE2 induced an expansion of reserve stem cells expressing *Ly6a, Clu, Msln*, and *Il1rn*, along with genes associated with regenerative and tumorigenic programs, via activation of *Ptger4* (EP4 receptor for PGE2 and YAP signaling pathways. All of these show that clusters 1-3 have a role in stem cell niche maintenance, not only through Wnt and Rspo secretion but also through PGE2. They are similar to Clusters 1 and 5 in our dataset.

### Myofibroblasts - clusters 4 and 5

Degirmenci et al. identified myofibroblasts as a cluster 2 (C2) population expressing *Acta2*, *Myh11* (smooth muscle myosin), and *Foxl1* (**Suppl. Figure 11**). While Degirmenci et al. focus on Foxl1 expression in myofibroblasts, they also find significant levels of *Vim* and *Sfrp1* expression in more than half of these cells, with *Gli1* and *Foxl1* expression in a smaller proportion of the total myofibroblast population. Kinchen et al. define myofibroblasts by their expression of *Aoc3*, nothing that *Foxl1* serves as a marker for both myofibroblasts and the Str2 subpopulation (telocytes) in their study. Degirmenci et al., Kinchen et al., and McCarthy et al. use *Myh11* to detect myofibroblasts, although we find that *Myh11* is characteristic rather for smooth muscle/pericytes cluster. Furthermore, Kinchen et al. suggest *Hhip* and *Pde6h* as potential markers. Roulis et al. take a more direct approach, identifying myofibroblasts based on high *Acta2* expression. They are similar to our Cluster 3.

### Smooth muscle (SM)/ pericyte-like fibroblasts-cluster 10

Kinchen et al. identifies this cluster by its high *Aoc3* expression, pointing out a gradient of *Acta2* with the highest expression in smooth musle cells and decreasing in myofibroblasts, pericytes (*Cspg4*^+^, known as *Ng2*) and telocytes (Str2) (**Suppl. Figure 12**). McCarthy et al. describe these cells based on the expression of *Rgs5* and *Pdgfrb*. This cluster corresponds closely to Cluster 4 in our dataset.

### Conclusion

This study, alongside recent reviews ^10–12,23^, illuminates the remarkable heterogeneity of subepithelial fibroblasts within the intestinal colonic mucosa. Using single-cell RNA sequencing analysis, immunofluorescence labeling, and imaging, we identified distinct fibroblast clusters with unique morphologies, localizations, and functions.

Our analysis indicates that consistent with previous observations, subepithelial fibroblasts express genes involved in ECM remodeling and cytokines that regulate the fate of epithelial cells, including stemness, proliferation, and differentiation. Surprisingly, these functions are distributed across different clusters, suggesting a specialization of roles. While trophocytes/ ECM fibroblasts and fibroblasts primarily produce extracellular matrix, myofibroblasts, and smooth muscle/pericyte-like fibroblasts are involved in contraction. This specialization challenges the previous belief that fibroblasts, in general, perform both functions simultaneously. Additionally, there is a functional division in the production of BMP/Wnt molecules. Telocytes contribute to BMP production, playing a role in epithelial differentiation, whereas trophocytes/ ECM fibroblasts and fibroblasts support the epithelial stem cell niche by producing Wnt molecules and BMP antagonists.

Moreover, subepithelial fibroblasts exhibit distinct shapes and localizations within the colonic mucosa but individual subtypes have similar morphology in the proximal and distal colons. Trophocytes/ ECM fibroblasts, fibroblasts, and telocytes are stellate-shaped and located close to the epithelium, forming a basket around the crypt. While trophocytes/ ECM fibroblasts produce ECM and cytokines essential for maintaining the stem cell niche, telocytes support the differentiation of epithelial cells.

In contrast, myofibroblasts and smoth muscle/pericyte-like fibroblasts are spindle-shaped. Some cells of these subpopulations span from the bottom to the top of the crypts, making contact with blood vessels on the crypt top. Further investigation into the function and development of this cluster is necessary, as close contact with the blood vessel may indicate a role in the control of blood vessel perfusion. Although the markers of contractile fibroblasts (myofibroblasts and SM/pericyte-like cells) are well described, the role of these cells in gut homeostasis remains unclear. Cell contractility is essential for physiological functions such as muscle contraction and cell migration, and it also plays a role in several human diseases, including fibrosis, scarring, and cancer. As described previously, besides muscle cells, myofibroblasts are another type of professional contractile cells. Typically arising from normal fibroblasts upon stress stimuli, activated contractile myofibroblasts (αSMA+) aid in wound repair by remodeling the connective tissue. Once the wound is repaired, myofibroblasts usually die. If they persist beyond the healing phase, they may contribute to scarring and fibrosis ^12^. Interestingly, unlike in other organs, contractile fibroblasts (αSMA+ myofibroblasts and SM/pericyte-like fibroblasts) are also present in gut homeostasis. Why does the gut need special contractile equipment of fibroblasts around its crypts? We hypothesize that they might be involved in the maintenance of mucosal architecture during feces transit or that they aid in mucus expulsion from the crypts, similar to myoepithelial cells, which aid in milk ejection from the breast glands ^28^.

The analysis of cell cycle markers revealed minimal proliferation within all fibroblast clusters except trophocytes/ ECM fibroblasts and fibroblasts, where a small subset of cells displayed slightly higher score levels. This suggests that most colonic fibroblasts do not proliferate in homeostasis, supported by Ki67 immunostaining. These results align with the previous studies suggesting that most fibroblasts reside in the G0/G1 phase of the cell cycle^7^. Trajectory analysis provided insights into the potential differentiation pathways of the colonic fibroblast subpopulations with major fibroblast subtypes (clusters) taking separate paths towards differentiation. The analysis also suggests that one of the fibroblast clusters could contain progenitors, but this warrants further investigation.

In summary, we propose a novel classification system for colonic intestinal fibroblasts. This system encompasses five distinct subpopulations: telocytes, trophocytes/ ECM fibroblasts, fibroblasts, myofibroblasts, and smooth muscle (SM)/ pericyte-like fibroblasts. All characteristics of different fibroblast subpopulations are summarized in the table below (**Table 1**). Signatures of different fibroblast populations are projected onto UMAP plots using AddModuleScore function (**Suppl. figure 6**).

## Acknowledgments

We thank all members of the Vignjevic and Lennon labs for helpful discussions. We also thank J. Barbazan and D. Krndija for help with the mouse experiment. We acknowledge Vincent Fraisier, Chloe Guedj and Lucie Sengmanivong from the Cell and Tissue Imaging Facility (PICT-IBiSA). This work received funding from European Research Council (ERC), under the grant agreement CoG 772487, ANR FIBROWOUND (DMV). DMV is an INSERM researcher.

## Material and methods

### Mice

Mice animal care and use for this study were performed in accordance with the European and French National Regulations for the Protection of Vertebrate Animals used for Experimental and other Scientific Purposes (2010/63/UE) for the care and use of laboratory animals. All experimental procedures were approved by the ethics committee of the Institut Curie CEEA-IC #118 amendment to the project 2018-004 (Autorisation: APAFIS #18510-2019011614361196 v1) given by National Authority in compliance with the international guidelines.

All mice were maintained in the Institut Curie (Paris, France) SPF animal facility before use. αSMA:Cre-ER^T2^/R26^mT/mG^ mouse was created in the host laboratory by crossing R26^mT/mG^ with a mouse expressing Cre-ER^T2^ recombinase under the αSMA promoter. Induction of Cre-ER^T2^ in animals was carried out through intraperitoneal injections of tamoxifen dissolved in sunflower seed oil (55.5mg/kg per one injection/day) for 5 consecutive days. Upon tamoxifen injections, cell membranes of cells with active αSMA promotor became green. All the other cell membranes remained red (Tomato+). C57BL/6J mice were purchased from Charles River (controls for the unmixing procedure, see later). Experiments were performed on 8 to 20 weeks old male or female mice.

### Tissue preparation

To achieve the same segmentation of the colons across the experiments, the intestine was removed from the animals, and washed with DPBS (MgCl_2_^-^, CaCl_2_^-^) at room temperature (RT). Different segments of the intestine were processed as follows: the basin was cut open at the pubic symphysis, anus and rectum were cut out (approximately 4 mm from the anus), the proximal colon was defined as the first quarter of the residual tube, starting from the caecum, while the distal colon was defined as the last quarter of the colon. Different segments were separated with the scalpel and further cut perpendicularly to the intestinal length into 3-4 mm pieces. The tissue was fixed in 4% Paraformaldehyde (PFA) in DPBS (MgCl_2_^+^, CaCl_2_^+^), 5 mL per segment, for 1-3h at RT with mild agitation. PFA 4% solution was prepared on the same day with 16% PFA stock solution (EMS, 15710), 10xDPBS (MgCl_2_^+^, CaCl_2_^+^, 1x final concentration), and distilled water. Upon fixation, tissue was washed three times in DPBS (MgCl_2_^-^, CaCl_2_^-^) and processed for staining.

### General whole-mount staining procedure

Fixed tissue (3-4 mm pieces) was cut perpendicularly to the intestinal length into thin slices either with a scalpel (0.5-1mm) or vibratome (0.3mm), well described previously ^29^, and the slices were placed into the Eppendorf tubes filled with DPBS (MgCl_2_^-^, CaCl_2_^-^) (1 tube per staining, 4-6 sections per tube). While cutting on the vibrotome, we adjusted the speed to 1.5mm/s, amplitude 0.7-0.9mm, and we used 3% agarose in DPBS solution.

Tissue was permeabilized with 1% Triton X-100 in DPBS (MgCl_2_^-^, CaCl_2_^-^) for 1h at room temperature (RT) with constant agitation. Blocking was done in 0.2% Triton X-100 in DPBS (MgCl_2_^-^, CaCl_2_^-^) solution containing 3% Bovine Serum Albumin (BSA) and goat or donkey serum (1:20 dilution, depending on the host of origin of secondary antibodies), for 1h at RT. Upon blocking, tissue slices were incubated with primary antibodies overnight at RT. Afterward, the tissue was washed three times in 0.2% Triton X-100 in DPBS (MgCl_2_^-^, CaCl_2_^-^) for up to 30 min/wash, incubated with secondary antibodies (and DAPI, depending on staining) for 2-4 h at RT, washed three times in 0.2% Triton X-100 in DPBS (MgCl_2_^-^, CaCl_2_^-^) for up to 1h/wash and mounted on slides using AquaPolyMount. Slides were left to dry with coverslips facing downwards, causing the tissue to “sink” closer to the coverslip, thereby facilitating easier imaging later. In summary, all washes were done in 0.2% Triton X-100 in DPBS (MgCl_2_^-^, CaCl_2_^-^) with constant agitation, while incubations with blocking and antibodies were done without agitation (120 μl of antibodies or blocking solution/ Eppendorf tube).

### Diverse staining combinations, confocal microscopy, image analysis and processing

Images were acquired using inverted confocal microscopes (Leica DMi8, SP8 scanning head unit), equipped with HC PL APO CS2 40x/1.30 OIL objective, with a regular pixel size of 284 nm (x, y dimensions), 0,5 −1 um (z dimension) and a resolution of 1024×1024 or 2048×2048 pixels. For all types of acquisition, Hybrid Detectors were used. Image acquisition and microscope control were performed using the Leica LAS X software.

Image processing and analysis were done using Leica LAS X, Fiji, Imaris, or Imaris Viewer software. All images were corrected for brightness and contrast. Clipping plane, ortho slicer, oblique slicer, as well as sectioning option, were used to choose a specific plane of view in Imaris Viewer.

To spatially map different fibroblast subpopulations and visualize them simultaneously, together with nuclei and actively dividing cells (Ki67+), we developed three staining combinations (**Table 2**) comprising five or six colors.

**Table 2.**
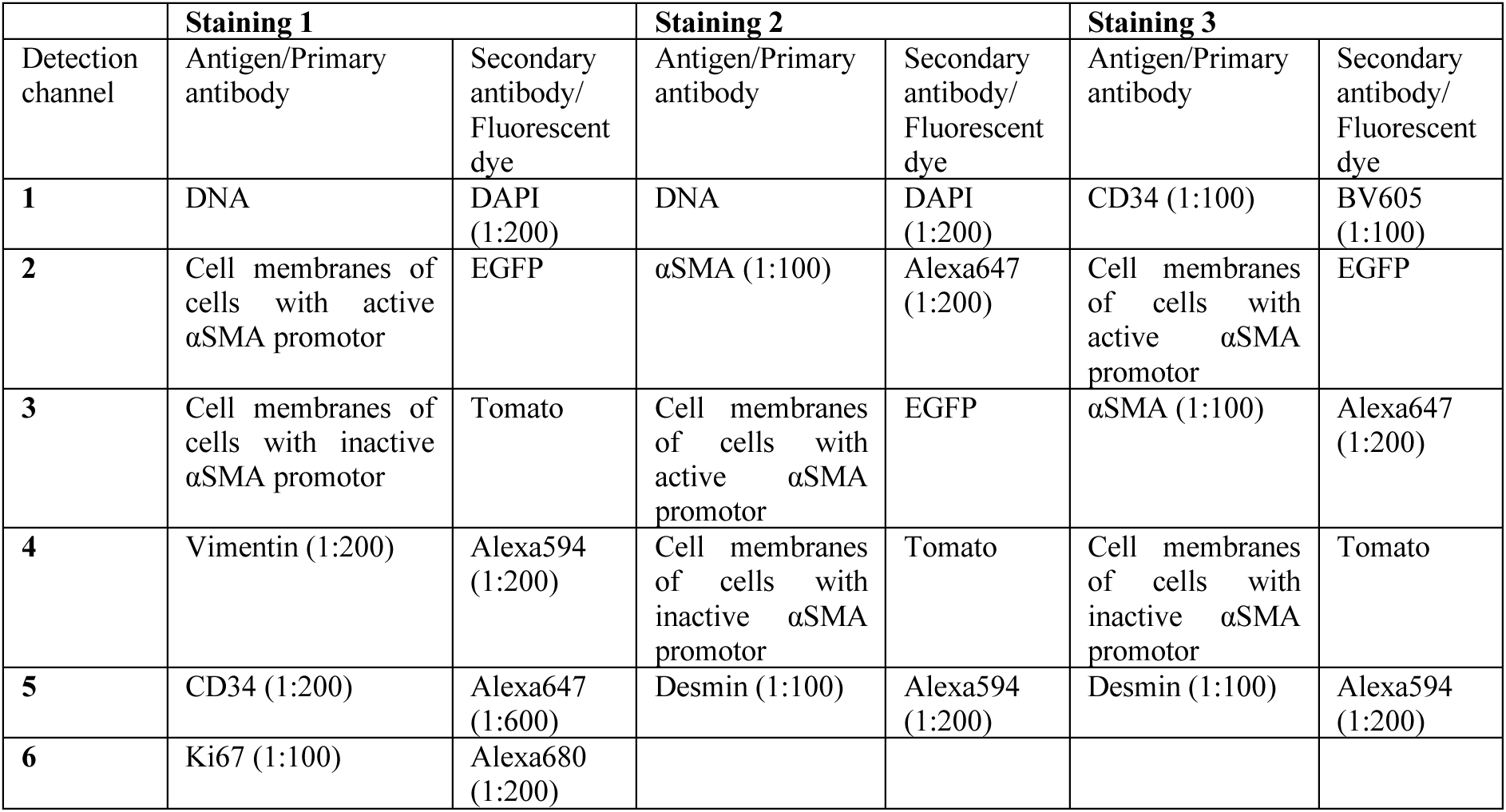
Staining combinations.

|  | Staining 1 |  | Staining 2 |  | Staining 3 |  |
| --- | --- | --- | --- | --- | --- | --- |
| Detection channel | Antigen/Primary antibody | Secondary antibody/ Fluorescent dye | Antigen/Primary antibody | Secondary antibody/ Fluorescent dye | Antigen/Primary antibody | Secondary antibody/ Fluorescent dye |
| 1 | DNA | DAPI (1:200) | DNA | DAPI (1:200) | CD34 (1:100) | BV605 (1:100) |
| 2 | Cell membranes of cells with active αSMA promotor | EGFP | αSMA (1:100) | Alexa647 (1:200) | Cell membranes of cells with active αSMA promotor | EGFP |
| 3 | Cell membranes of cells with inactive αSMA promotor | Tomato | Cell membranes of cells with active αSMA promotor | EGFP | αSMA (1:100) | Alexa647 (1:200) |
| 4 | Vimentin (1:200) | Alexa594 (1:200) | Cell membranes of cells with inactive αSMA promotor | Tomato | Cell membranes of cells with inactive αSMA promotor | Tomato |
| 5 | CD34 (1:200) | Alexa647 (1:600) | Desmin (1:100) | Alexa594 (1:200) | Desmin (1:100) | Alexa594 (1:200) |
| 6 | Ki67 (1:100) | Alexa680 (1:200) |  |  |  |  |

In these staining combinations employing more than four colors, an unmixing approach was used to remove fluorophores spillover. As described previously^30^, we used single-stained tissues to obtain compensation coefficients that were applied to perform spectral unmixing using a homemade Fiji macro. Single stained tissues were tissue slices obtained from non-treated non-fluorescent C57BL/6J mice and stained just for one particular antigen using the same concentrations of antibodies as the one used in a full panel of antibodies, following the same experimental procedure already described. Single color controls of αSMA:Cre-ER^T2^/R26^mT/mG^ mouse (Tomato and EGFP) were non-stained tissue slices obtained from it. Stainings that label the same cell structures were put in channels with minimized cross-talk. To obtain compensation coefficients, single stained tissues were imaged with the same acquisition settings used for tissues stained with the whole set of antibodies. Coefficients were calculated as described previously and later used for spectral unmixing using spectral-unmixing macro code in Fiji (https://doi.org/10.17632/bdfyprfsv9.1) ^30^. More information about the staining combinations are given in the **Table 2**.

All other staining combinations consisted of four colors, including Tomato and EGFP, which are inherent mouse fluorophores, and two antigens stained with secondary antibodies Alexa405 and Alexa647. References of all primary and secondary antibodies used are given in the table at the end of the document.

### Isolation of intestinal cell suspensions and flow cytometry

Five αSMA:Cre-ER^T2^/R26^mT/mG^ mice have been injected for 5 consecutive days with Tamoxifen (50 mg/kg) and killed 2 days after. After colon dissection, tissue was washed 3 times with ice-cold L15 media (MgCl_2_^-^, CaCl_2_^-^) and cut open longitudinally. Then, mucosal and muscle layers were separated with the help of glass-slide scraping. The mucosal sample was mechanically dissociated using 2 scalpels and digested with collagenase from Clostridium histolyticum solution (C2139, Sigma, 100U/mL), enriched with growth factors (DMEM-Glutamax with 0.25 U/mL insulin, 100 µg/mL transferrin, 10 ng/mL murine EGF and 1% antibiotic-antimycotic) and incubated for 1h at 37°C with constant agitation with magnetic stirrer. EGFP + cells were enriched by FACS sorting on BD Aria II sorter and processed with FlowJo vX software.

### Single Cell RNA-sequencing using dropseq

#### RNaseq libraries preparation

Cellular suspension (∼5000 cells, with expected recovery of ∼3000 cells) of sorted fibroblast from the entire colon of 10 weeks old αSMA:Cre-ER^T^^2^/R26^mT/mG^ mice (5 mice in total) was loaded on the 10X Chromium Controller instrument (10X Genomics) according to the manufacturer’s protocol, based on the 10X GEMCode proprietary technology.

The Chromium Single Cell 3’ Library Kit V3 (10X Genomics) was used to generate the cDNA and prepare the libraries, according to the manufacturer’s protocol. The libraries were then equimolarly pooled and sequenced on a Novaseq 6000 (Illumina). A coverage of 400M reads per sample was targeted, in order to obtain 100 000 reads per cell. The raw data were then demultiplexed and processed with the Cell Ranger software Version 2.0.1 and 3.0.1 on fastq files (10X Genomics).

### Quality check, read alignment, and computation of the UMI counts scRNA-seq pre-processing

Raw data is deposited on the GEO under the reference GSE269017 (https://www.ncbi.nlm.nih.gov/geo/query/acc.cgi?acc=GSE269017). During the revisions, the private token to access the data is: **srwtiqcetvylxwz**.

The scRNA-Seq data analyses are performed by GenoSplice technology (www.genosplice.com). Sequencing data quality analysis was performed using FastQC v0.11.2. For read alignment and unique molecular identifiers (UMI) quantification, CellRanger software v3.0.2 was used on Mouse mm10 genome, with Human EGFP sequence added. Default parameters were applied, except for using Mouse FAST DB 2018_1 annotations and Human EGFP sequence. The expression matrix containing UMI counts was filtered to retain only genes with UMI ≥ 1 in at least one cell. Two subdatasets were generated: a “fibroblast” dataset with cells expressing Vimentin, not expressing H2-Ab1, and with mitochondrial expression ≤20%, and an “epithelial” dataset with cells not expressing Vimentin and expressing Keratin 8. For UMIs normalization, Seurat 2.3.4 was used [PMID:29608179], and global-scaling normalization method was applied with a scale factor of 10000 and log-transformation of data. This was followed by a scaling linear transformation step, to avoid highly-expressed genes having higher weight in downstream analysis.

### Clustering and marker genes

PCA was performed on the scaled data, with a Jackstraw plot to choose how many PCs to retain as an input for Seurat clustering step (UMAP on the Figure 1 was created using PC=20). Default parameters were used, with Louvain algorithm as the clustering method and a resolution parameter defining the clusters granularity set to 0.6. Marker genes defining each cluster were found via differential expression testing, with a log fold change threshold of 1.

### Trajectory

Single-cell trajectories were constructed using Monocle 3 ^31–34^ as well as Slingshot 2.10 ^35^. In both analyses, cluster 1 was predefined as trajectories’ root.

For RNA Velocity^36^ spliced and unspliced expression matrices were generated using the standard velocyto pipeline for Epi and LP samples. Loom files were merged using loompy package on Python. R packages velocyto.R and SeuratWrappers were then used to estimate RNA velocity vectors with velocity parameters kCells = 25, fit.quantile = 0.2 and deltaT = 1 and visualization parameters n = 200, grid.n = 40, arrow.scale = 3 and scale = “sqrt.”

### Integration with public datasets

Datasets GEO GSE113043 ^6^ (Degirmenci dataset: 1 sample), GEO GSE114374 ^7^ (Kinchen dataset: 3 samples), GEO GSE130681^8^ (McCarthy dataset: 2 samples) and GEO GSE142431^9^ (Roulis dataset: 5 samples) have been integrated and compared with our dataset.

In order to compare cluster definitions, each dataset has been reanalyzed using the publication thresholds and parameters, finally by comparing markers, we redefined clusters from the original analysis. After suppressing immune clusters from Kinchen and McCarthy datasets using *Ptprc* expression as a marker, all datasets were integrated with Seurat 3.2 CCA algorithm.

This Integrated dataset was clustered in the same way as previously, using 15 PCs and a resolution parameter of 0.8, defining 19 clusters.

### G2M/S score

G2M/S score is accessed as described previously ^37^. The genes corresponding to each part of the cell cycle are listed below.

S phase genes:

*Mcm5,Pcna,Tyms,Fen1,Mcm2,Mcm4,Rrm1,Ung,Gins2,Mcm6,Cdca7,Dtl,Prim1,Uhrf1,Mlf1ip,Hells,Rf c2,Rpa2,Nasp,Rad51ap1,Gmnn,Wdr76,Slbp,Ccne2,Ubr7,Pold3,Msh2,Atad2,Rad51,Rrm2,Cdc45,Cdc 6,Exo1,Tipin,Dscc1,Blm,Casp8ap2,Usp1,Clspn,Pola1,Chaf1b,Brip1,E2f8 Mlf1ip* is not detected in Glisovic, but detected in integrated data.

G2/M phase genes

*Hmgb2,Cdk1,Nusap1,Ube2c,Birc5,Tpx2,Top2a,Ndc80,Cks2,Nuf2,Cks1b,Mki67,Tmpo,Cenpf,Tacc3,F am64a,Smc4,Ccnb2,Ckap2l,Ckap2,Aurkb,Bub1,Kif11,Anp32e,Tubb4b,Gtse1,Kif20b,Hjurp,Cdca3,Hn 1,Cdc20,Ttk,Cdc25c,Kif2c,Rangap1,Ncapd2,Dlgap5,Cdca2,Cdca8,Ect2,Kif23,Hmmr,Aurka,Psrc1,An ln,Lbr,Ckap5,Cenpe,Ctcf,Nek2,G2e3,Gas2l3,Cbx5,Cenpa*

### G profiler enrichment analysis

The g: Profiler ^17^, an extensive webserver dedicated to functional enrichment analysis, was used for the gene enrichment analysis. Specifically, we employed g:GOSt within this platform, which compares a custom list of genes provided by the user against various established functional information sources, including Gene Ontology (GO), Kyoto Encyclopedia of Genes and Genomes (KEGG), Reactome, and WikiPathways (WP) (see **Suppl. Table 1** for the analysis links and all the parameters used). We have identified statistically significant biological processes, pathways, regulatory motifs, and protein complexes (Raudvere et al., 2019). For this analysis, we have generated gene lists for each cluster, using Seurat v 5.0.1, setting *log.fc* of gene expression to 0.5, and selecting only up-regulated genes in each cluster (**Suppl. Table 1).**

### Data visualization

Data visualization (DotPlots, FeaturePlots, ViolinPlots, HeatMap) was done using Seurat 5.0.1, along with the package “scCustomize” ^38^. The list of genes used to generate the HeatMap in Figure 1 is available as **Suppl. Table 2**.

To visualize transcriptional signature of each of the clusters (**Suppl. Figure 6**), detailed in Table 1, we have used Seurat function “AddModuleScore”.

**A list of all the reagents used is given in Table 3**.

**Table 3.** Chemicals and antibodies.

| Reagent | Identifier | Source |
| --- | --- | --- |
| Desmin Polyclonal Antibody (Rabbit) | PA5-16705 | Invitrogen- Thermo Fisher Scientific |
| Monoclonal Anti-Actin, α-Smooth Muscle, clone 1A4, ascites fluid (Mouse) | A2547 | Sigma- Aldrich Chimie |
| CD34 Monoclonal Antibody (RAM34), eBioscience (Rat) | 14-0341-82 | Invitrogen- Thermo Fisher Scientific |
| DAPI (4', 6-diamidino-2-phenylindole, dihydrochloride) | D1306 | Invitrogen- Thermo Fisher Scientific |
| Anti-Vimentin antibody (Chicken) | ab24525 | Abcam |
| Anti-PDGFR alpha antibody [EPR22059-270] (Rabbit) | Ab203491 | Abcam |
| Anti- CD31 antibody (Rabbit) | Ab28364 | Abcam |
| Monoclonal Anti-Collagen, Type I antibody produced in mouse | C2456 | Sigma |
| Anti-Collagen Antibody, Type IV (Rabbit) | AB756P | Sigma |
| Anti-Laminin antibody produced in rabbit | L9393 | Sigma |
| Anti-Connexin-43 antibody produced in rabbit | C6219 | Sigma |
| Collagenase from Clostridium histolyticum | C2139 | Sigma |
| Bovine Serum Albumin (IgG-Free, Protease-Free) | 001-000-162 | Jackson ImmunoResearch |
| Normal Goat Serum | 005-000-121 | Jackson ImmunoResearch |
| Normal Donkey Serum | 017-000-121 | Jackson ImmunoResearch |
| DPBS (no Calcium, no Magnesium) | 14190-144 | ThermoFisher Scientific |
| 10xDPBS (Calcium, Magnesium) | 14080055 | ThermoFisher Scientific |
| Tritonx100 | T8787 | Sigma-Aldrich |
| PFA (Paraformaldehyde) | 30525-89-4 | Electron Microscopy Sciences |
| AquaPolyMount | 18606-5 | Polysciences |
| Tamoxifen | 0215673891 | MP Biomedicals |
| Sunflower seed oil | S5007 | Sigma-Aldrich |
| DMEM-GlutaMAX | 10566016 | ThermoFisher Scientific -Gibco |
| Antibiotic-antimycotic (anti-anti) | A5955 | Life Technologies |
| FBS (fetal bovine serum) | A3840001 | ThermoFisher Scientific |
| Insulin | / | Sigma-Aldrich |
| Transferrin | T4132 | Holo, Human Plasma, Calbiochem |
| EGF (Epidermal growth factor) | / | Peptrotech |
| Leibovitz's L-15 Medium | 11415064 | ThermoFisher Scientific |
| Goat anti-Chicken IgY (H+L) Secondary Antibody, Alexa Fluor 594 | A11042 | Invitrogen- Thermo Fisher Scientific |
| Brilliant Violet 605™ Goat anti-rat IgG (minimal x-reactivity) Antibody | 405430 | BioLegend |
| Goat anti-Rat IgG (H+L) Cross-Adsorbed Secondary Antibody, Alexa Fluor 647 | A21247 | Invitrogen- Thermo Fisher Scientific |
| Alexa Fluor® 680 AffiniPure Donkey Anti-Rabbit IgG (H+L) | 711-625-152 | Jackson ImmunoResearch |
| Donkey anti-Rabbit IgG (H+L) Highly Cross-Adsorbed Secondary Antibody, Alexa Fluor 594 | A21207 | Invitrogen- Thermo Fisher Scientific |
| F(ab')<sub>2</sub>-Goat anti-Mouse IgG (H+L) Cross-Adsorbed Secondary Antibody, Alexa Fluor 647 | A21237 | Invitrogen- Thermo Fisher Scientific |
| Donkey anti-Rabbit IgG (H+L) Highly Cross-Adsorbed Secondary Antibody, Alexa Fluor™ Plus 405 | A48258 | Invitrogen- Thermo Fisher Scientific |
| Goat anti-Rat IgG (H+L) Highly Cross-Adsorbed Secondary Antibody, Alexa Fluor™ Plus 405 | A48261 | Invitrogen- Thermo Fisher Scientific |
| Alexa Fluor Plus 647 Phalloidin | A30107 | Thermo Fisher Scientific |

