## Supplementary data for "Intestinal fibroblast heterogeneity: unifying RNA-seq studies and introducing consensus-driven nomenclature"

### Supplementary figures

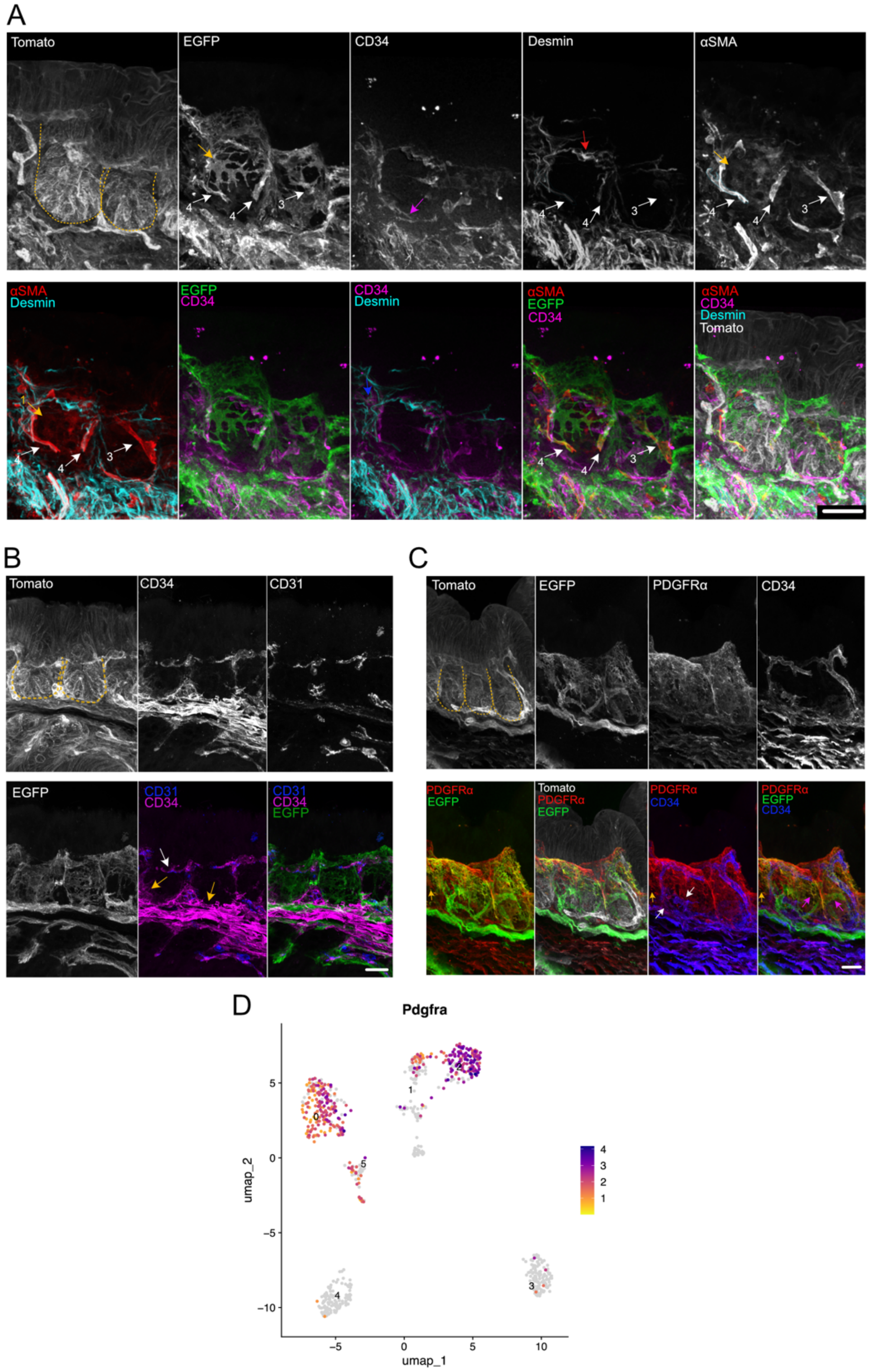


#### Supplementary Figure 1. Spatial organization of fibroblast populations in the proximal colon

Immunofluorescence of the proximal colon sections from αSMA:CreER$^{T2}$; R26$^{mT/mG}$ mice (scale bar, A: 40 µm, B and C: 20 µm). Cell membranes of αSMA+ fibroblasts are labeled with EGFP, all the other cell membranes are labeled with Tomato. Crypts are outlined with yellow dashed lines.

**A)** Tomato, EGFP, staining of desmin, αSMA protein, CD34. Yellow arrows mark stellate cells of cluster 2. They show strong EGFP expression, no CD34 and desmin expression, and low αSMA protein expression. White arrows, along with numbers 3 and 4, point to cells of clusters 3 and 4. While all three cells show high expression of EGFP and αSMA protein, the cells annotated with two numbers 4 have desmin, indicating they belong to cluster 4. Cell annotated with number 3 shows no staining of desmin, suggesting it is part of cluster 3. The pink arrow points to CD34+ cell (cluster 0 and 5).

**B)** Tomato, EGFP, staining of CD31, CD34. The white arrow points to the CD34+CD31+ blood vessel. Yellow arrows point to CD34+ fibroblasts (clusters 0 and 5).
**C)** Tomato, EGFP, staining of PDGFRα, CD34. The pink arrows point to EGFP+ PDGFRα + cells (cluster 2). The yellow arrow points to PDGFRα + cell (cluster 1) . White arrows show CD34+ PDGFRα+ cells (clusters 0 and 5).
**D)** Expression of *Pdgfra* is shown projected onto the UMAP plot.

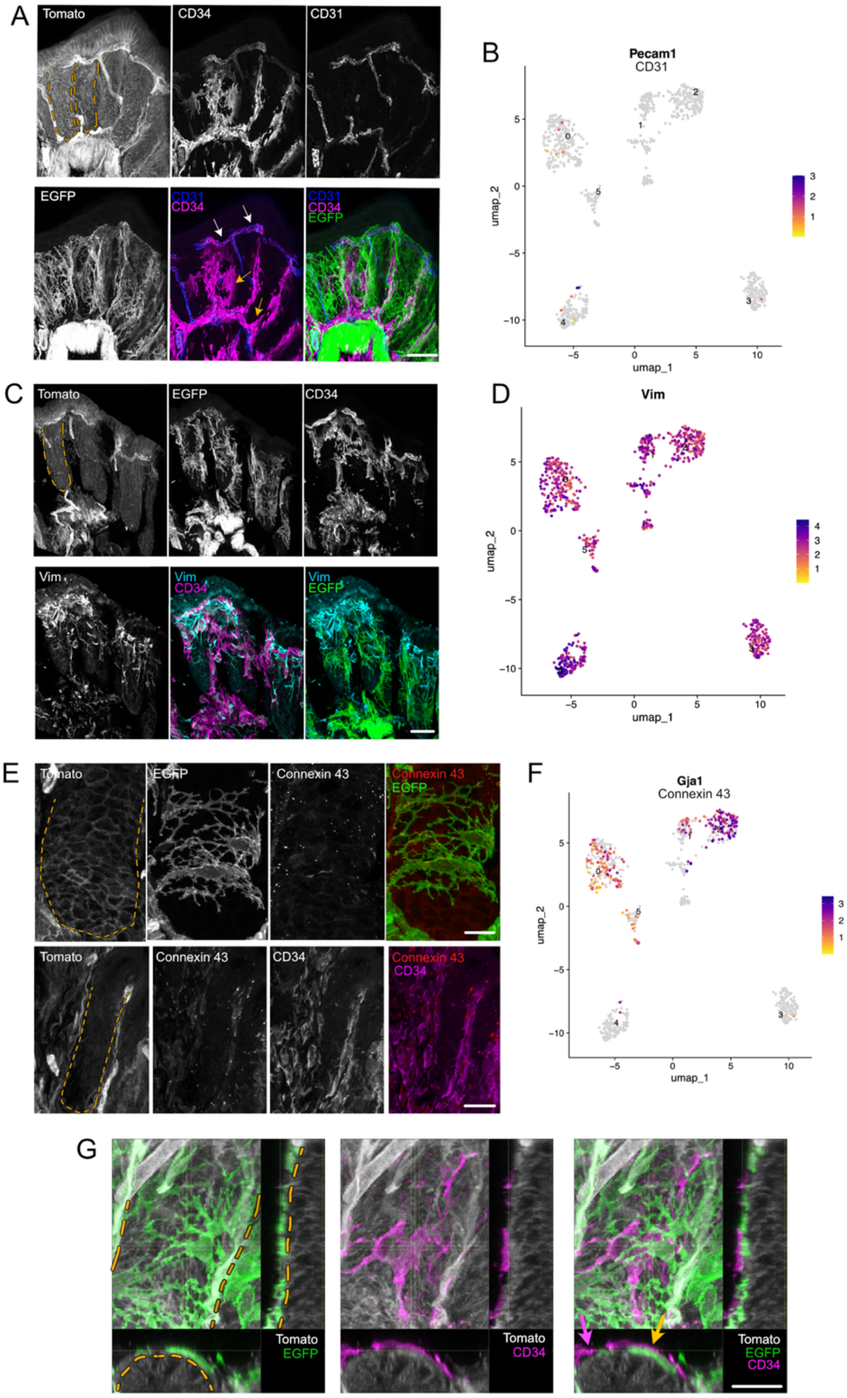


#### Supplementary Figure 2. Localization and gene expression patterns of fibroblasts in the distal colon

**(A, C, E)** Immunofluorescence of the distal colon sections from αSMA:CreER$^{T2}$; R26$^{mT/mG}$ mice (scale bar, A and C: 40 µm, E and G: 20 µm). Cell membranes of αSMA+ fibroblasts are labeled with EGFP, all the other cell membranes are labeled with Tomato. Crypts are outlined with yellow dashed lines.
**A)** Tomato, EGFP, staining of CD31, CD34. White arrows point to the CD34+CD31+ blood vessel. Yellow arrows point to CD34+ fibroblasts (clusters 0 and 5).
**C)** Tomato, EGFP, staining of Vimentin and CD34.

**E)** Tomato, EGFP, staining of Connexin43 and CD34.
**G)** Tomato, EGFP, staining of CD34. Different projections of the crypt - xy, xz, yz. αSMA+ fibroblasts labeled with EGFP are in direct contact with crypt epithelium (Tomato). CD34+ fibroblasts are on top of αSMA+ fibroblasts (EGFP+; yellow arrow). Pink arrow points to CD34+ fibroblasts which are in direct contact with crypt epithelium (Tomato).
**B)** Expression of CD31 (*Pecam1*) is shown projected onto the UMAP plot.
**D)** Expression of Vimentin (*Vim*) is shown projected onto the UMAP plot.
**F)** Expression of Connexin43 (*Gja1*) is shown projected onto the UMAP plot.

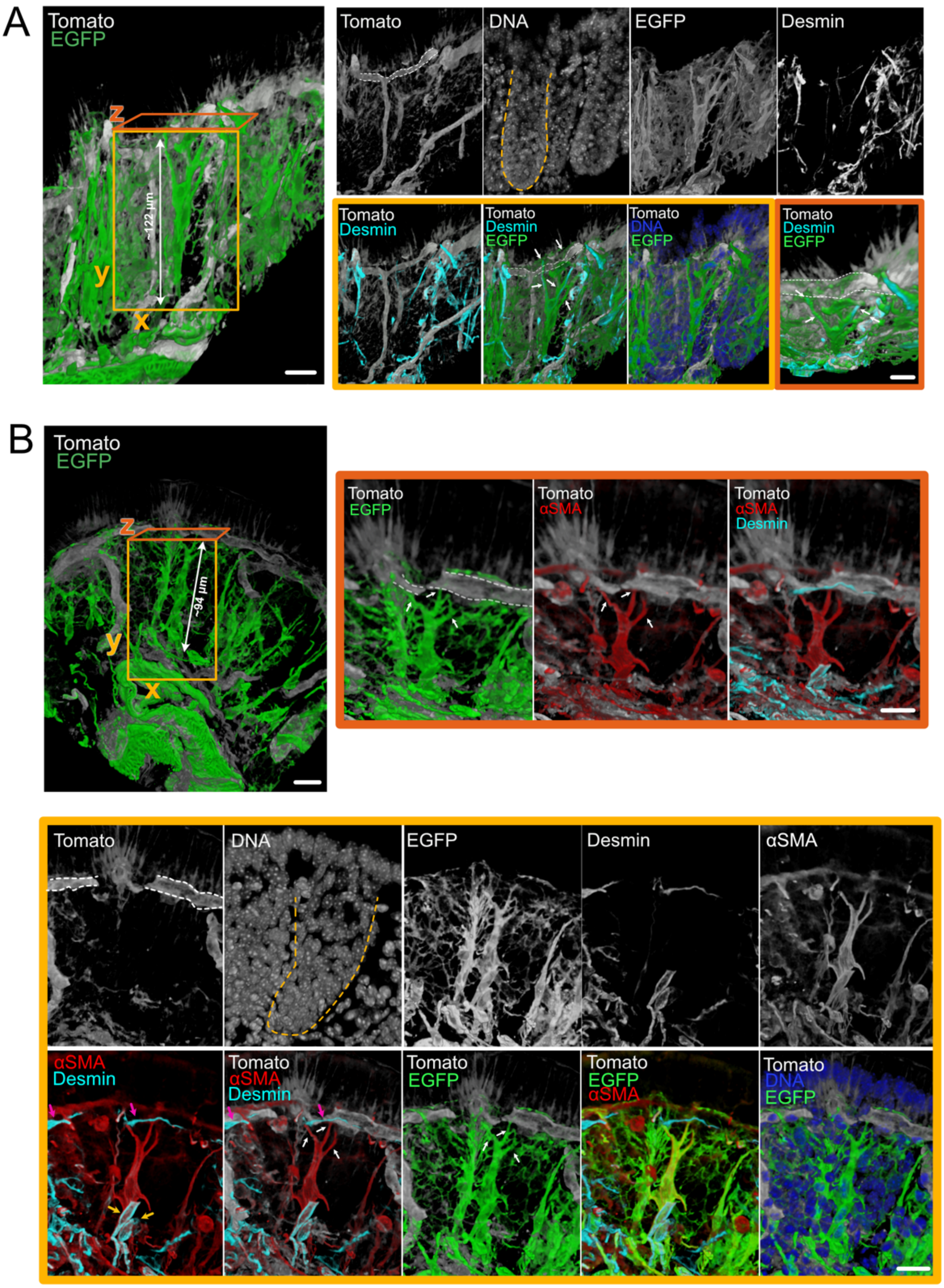


#### Supplementary Figure 3. Spatial organization of spindle and stellate fibroblasts in the distal colon

Immunofluorescence of the distal colon sections from αSMA:CreER$^{T2}$; R26$^{mT/mG}$ mice (scale bar, 20 µm). Cell membranes of αSMA+ fibroblasts are labelled with EGFP, all the other cell membranes are labelled with Tomato. Crypts are outlined with yellow dashed lines, and blood vessels with white dashed lines. Images are made in the blend mode using Imaris Viewer.

**A)** Tomato, EGFP, staining of DNA, and desmin. On the left-hand side, big field of view, xy plane. On the right, first row and the second row- in the yellow square: zoom-in on the long stellate fibroblast, xy view, in the orange square: xz view. White arrows point to the long “arms” of the spindle cell (cluster 4, since it expresses desmin) touching the blood vessel.
**B)** Tomato, EGFP, staining of DNA, desmin and αSMA. In the first row, on the left-hand side, big field of view, xy plane. In the same row, on the right, in the orange square, xz view of the spindle fibroblast. White arrows point to the long “arms” of the spindle cell (cluster 3, since it expresses αSMA, but not desmin), touching the blood vessel. In the second and third row, in the yellow square, zoom-in on the long stellate fibroblast, xy view. Yellow arrows point to the cells of the cluster 4 (expressing αSMA and desmin, connecting to the muscle layer below them). Pink arrows point to desmin+ cells on top of the blood vessels, most likely pericytes.

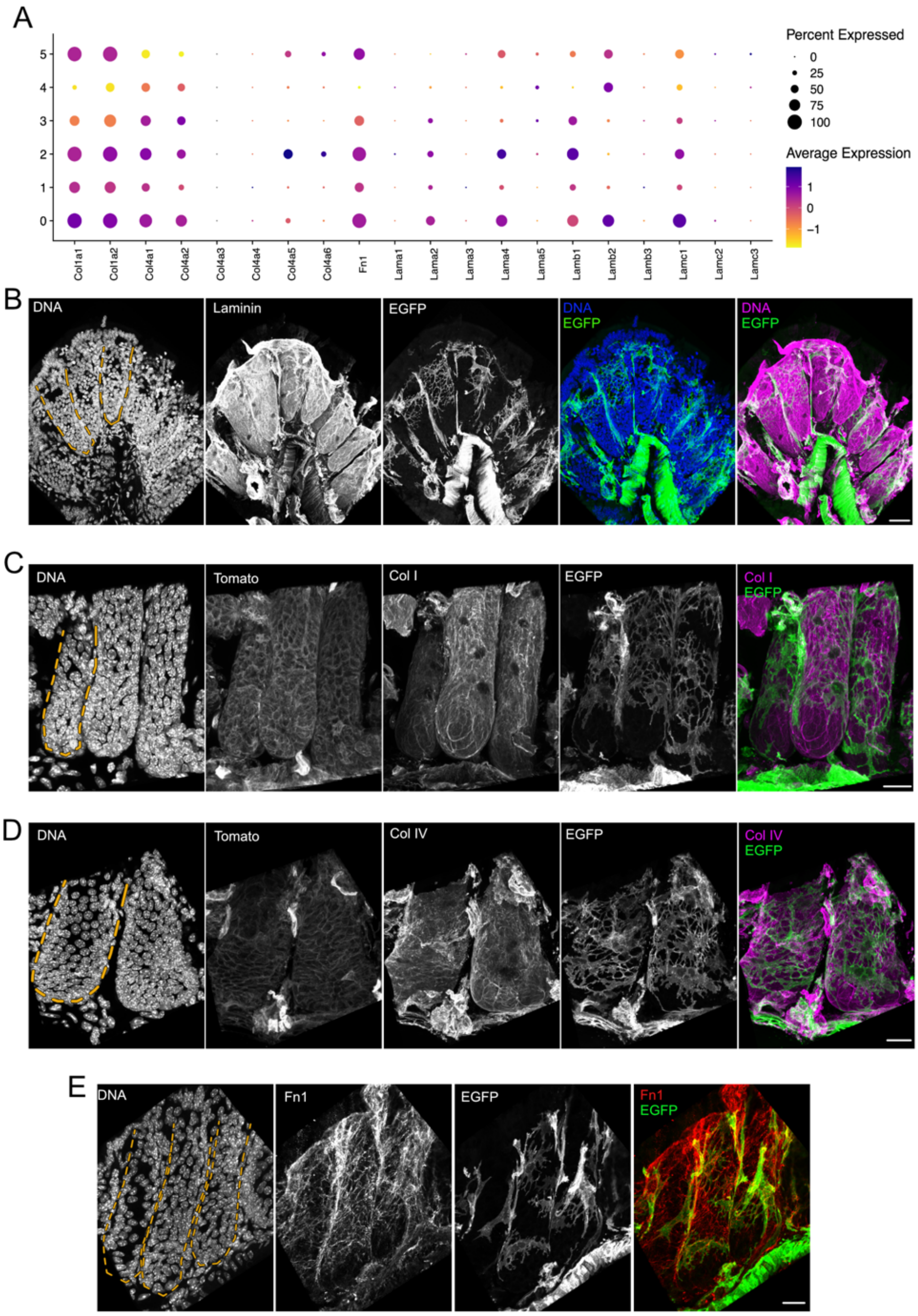


#### Supplementary Figure 4. ECM gene expression and protein localization in the distal colon

**A)** Dot plot of the relative expression of some ECM genes (collagens, fibronectin, and laminins). The size of the dot represents the percentage of cells that express the transcript, while the color of the dot is the average expression level within a cluster.

**(B-E)** Immunofluorescence of the distal colon sections from αSMA:CreER$^{T2}$; R26$^{mT/mG}$ mice (scale bar, 20 µm). Cell membranes of αSMA+ fibroblasts are labeled with EGFP, all the other cell membranes are labeled with Tomato. Crypts are outlined with yellow dashed lines.
**B)** Tomato, EGFP, staining of DNA, and laminin.
**C)** Tomato, EGFP, staining of DNA, and collagen I.
**D)** Tomato, EGFP, staining of DNA, and collagen IV.
**E)** Tomato, EGFP, staining of FN1.

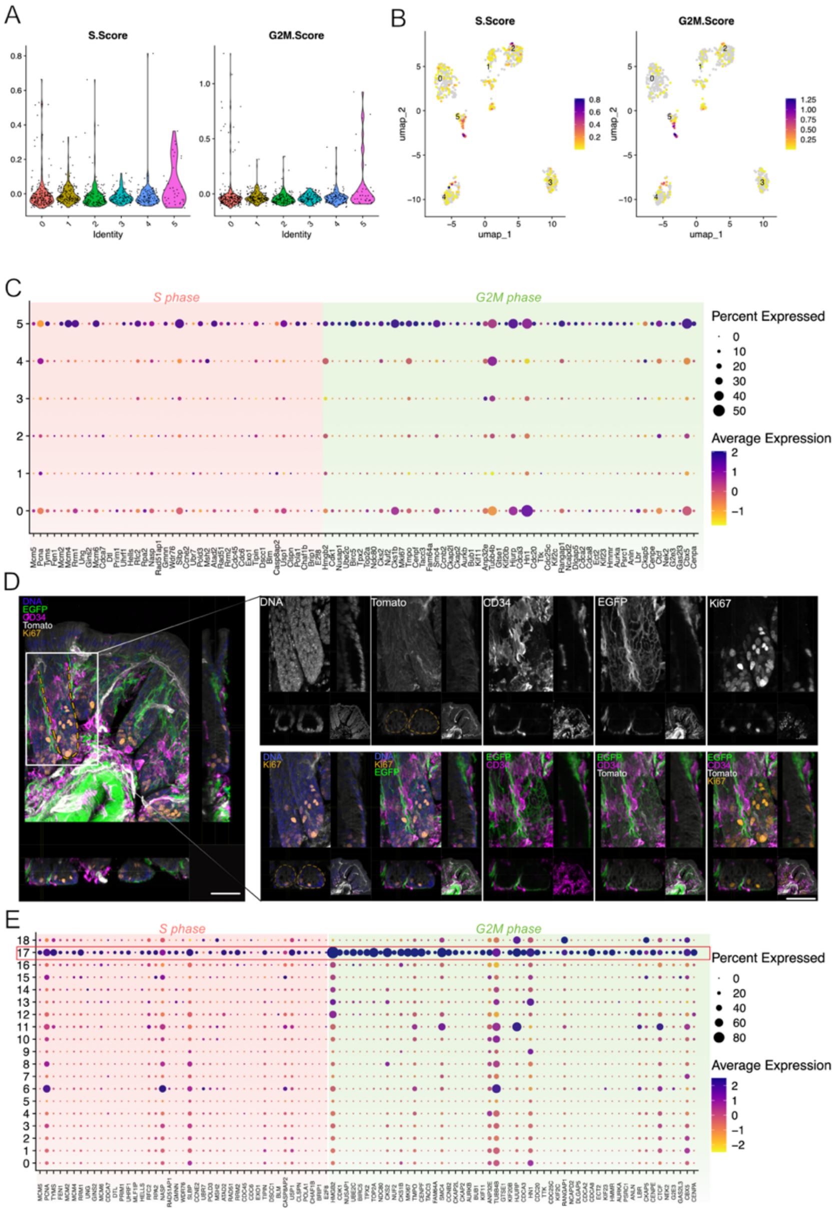


#### Supplementary Figure 5. Cell cycle analysis of fibroblasts

**A)** Violin plots showing S.Score and G2M.Score for each fibroblast cluster.
**B)** S.Score and G2M.Score are shown projected onto UMAP plots.
**C)** Dot plot illustrating the relative expression levels of marker genes associated with the G2/M and S phases of the cell cycle (Glisovic dataset). The size of the dot represents the percentage of cells that express the transcript, while the color of the dot is the average expression level within a cluster.
**D)** Immunofluorescence of the distal colon sections from αSMA:CreER$^{T2}$; R26$^{mT/mG}$ mice (scale bar, 40 µm). Cell membranes of αSMA+ fibroblasts are labeled with EGFP, all the other cell membranes are labeled with Tomato. Crypts are outlined with yellow dashed lines. On the left side, Tomato, EGFP, staining of DNA, CD34, and Ki67. Colonic crypts are shown in various projections (xy, xz, yz). On the right side, a closer view of one of the crypts in xy, xz, and yz projections. Notably, there are no Ki67+ fibroblasts observed, neither of αSMA+ fibroblasts are labeled with EGFP, nor CD34+ fibroblasts.

**E)** Dot plot illustrating the relative expression levels of marker genes associated with the G2/M and S phases of the cell cycle (Merged/Integrated dataset). The size of the dot represents the percentage of cells that express the transcript, while the color of the dot is the average expression level within a cluster. Cluster 17 shows the highest expression of G2/M and S markers.

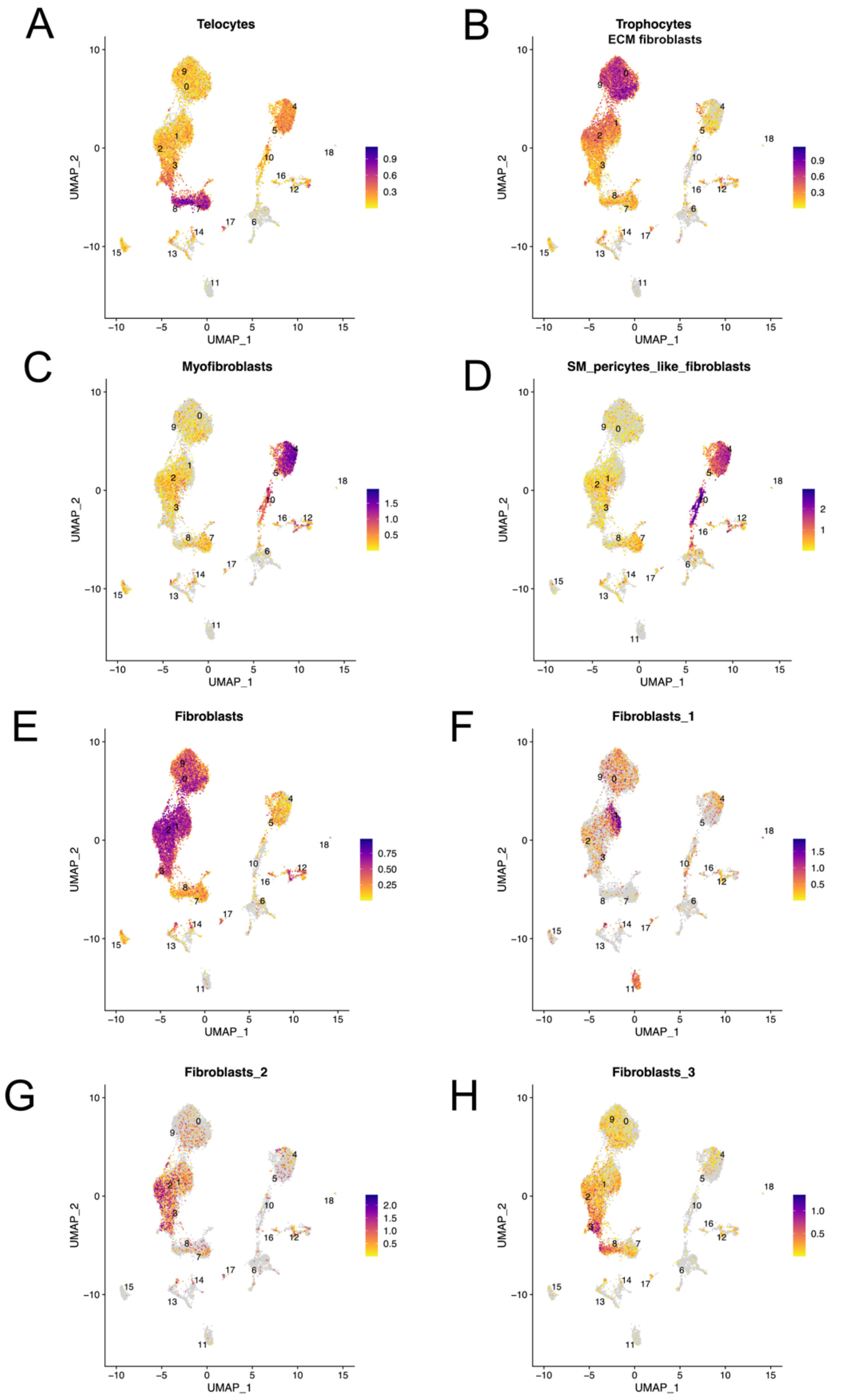


**Supplementary Figure 6. UMAP visualization of fibroblast population signatures**

**(A-H)** Signatures of different fibroblast population projected onto UMAP plots using AddModuleScore function.

### Telocytes

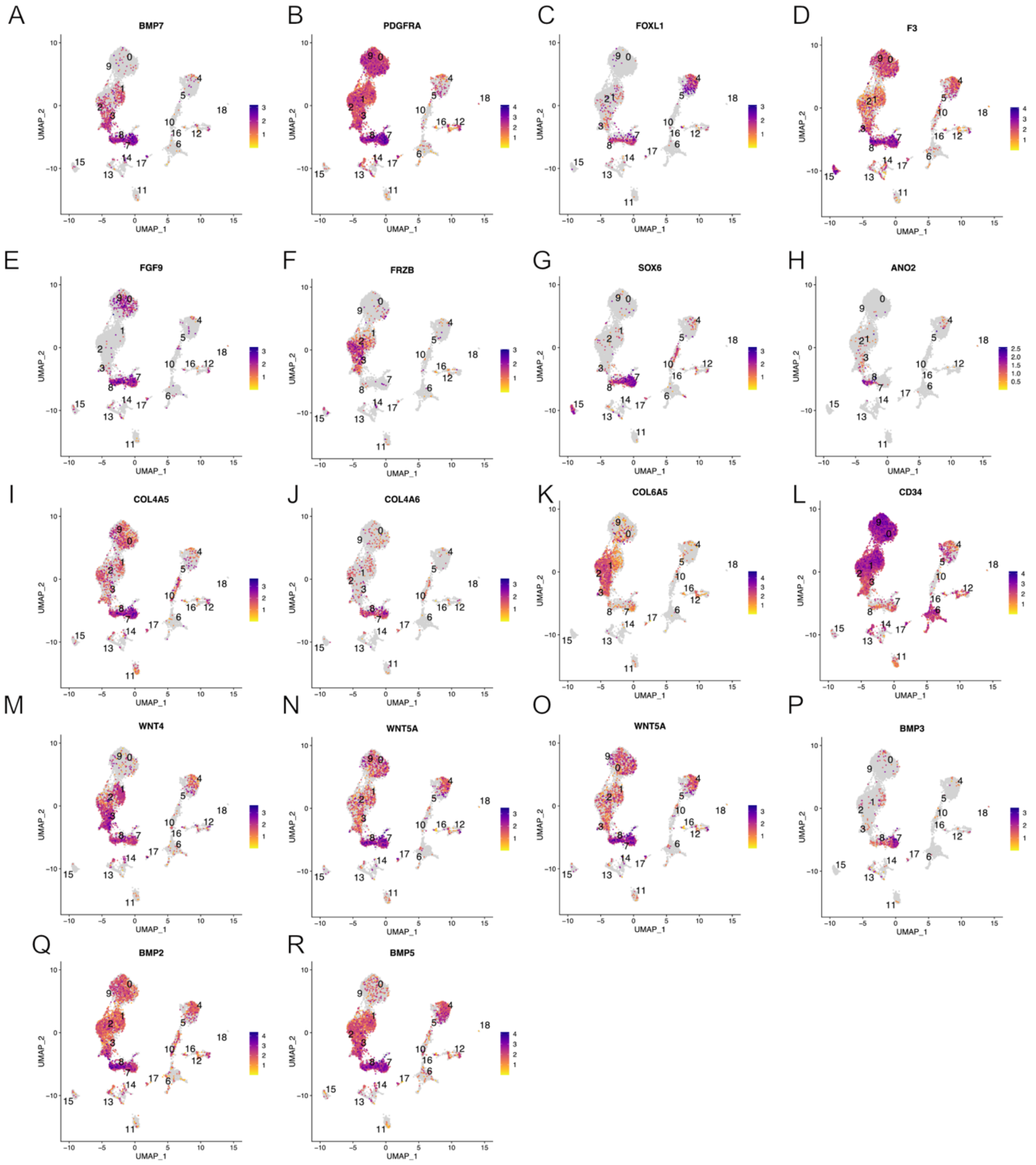


**Supplementary figure 7. Telocytes**

**(A-R)** Expression of different telocyte markers projected onto UMAP plots.

### Trophocytes/ ECM fibroblasts

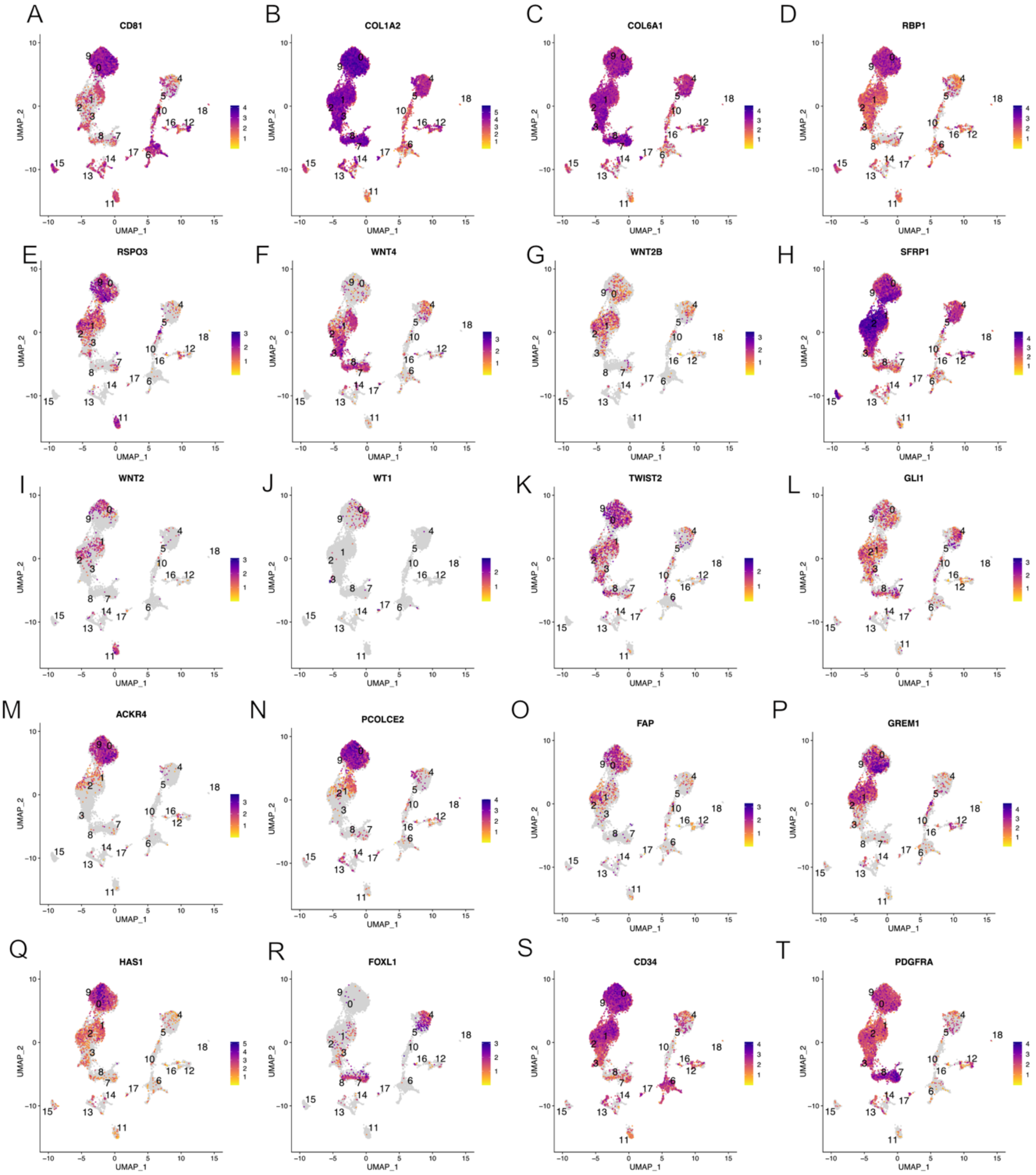


**Supplementary figure 8. Trophocytes/ ECM fibroblasts**

**(A-T)** Expression of different trophocyte/ECM fibroblasts markers projected onto UMAP plots.

### Trophocytes/ ECM fibroblasts

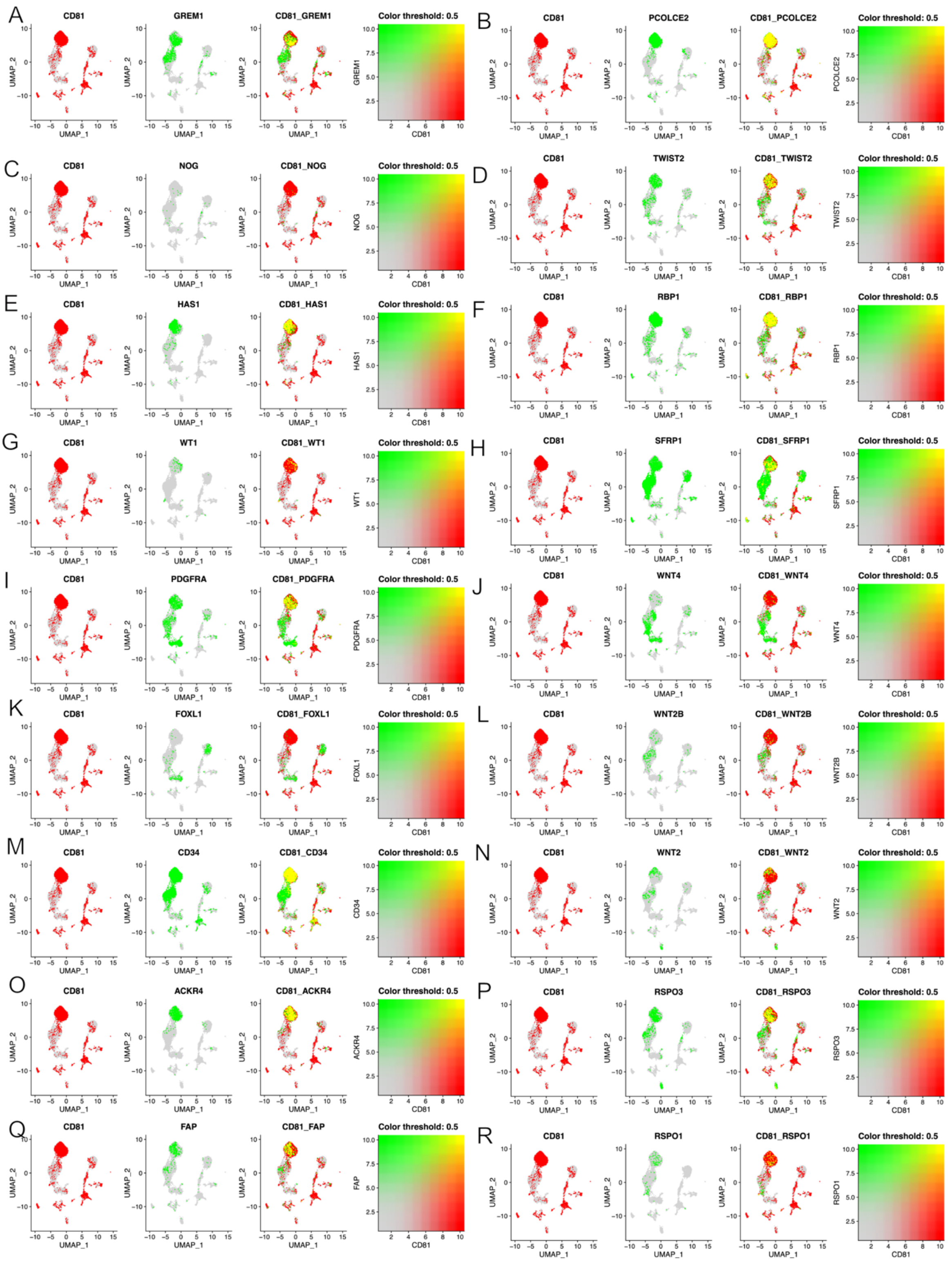


**Supplementary figure 9. Trophocytes/ ECM fibroblasts**

**(A-R)** Visualization of two marker genes of trophocytes/ ECM fibroblasts projected onto UMAP plots one at a time, then simultaneously, with overlapping scores.

### Fibroblasts- Clusters 1, 2, 3

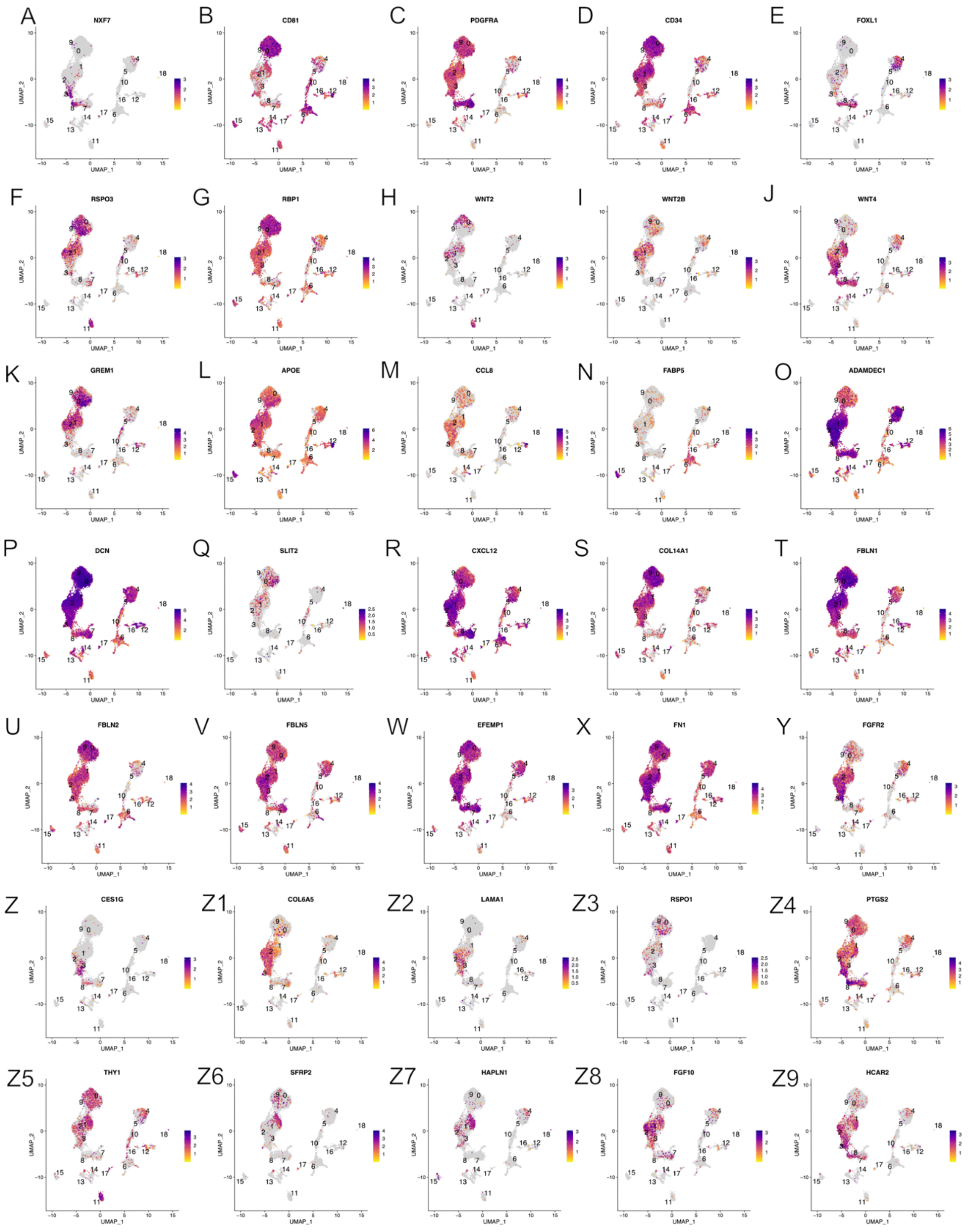


**Supplementary figure 10. Fibroblasts**

**(A-Z, Z1-Z9)** Expression of different markers of fibroblast (clusters 1, 2, 3) projected onto UMAP plots.

### Myofibroblasts

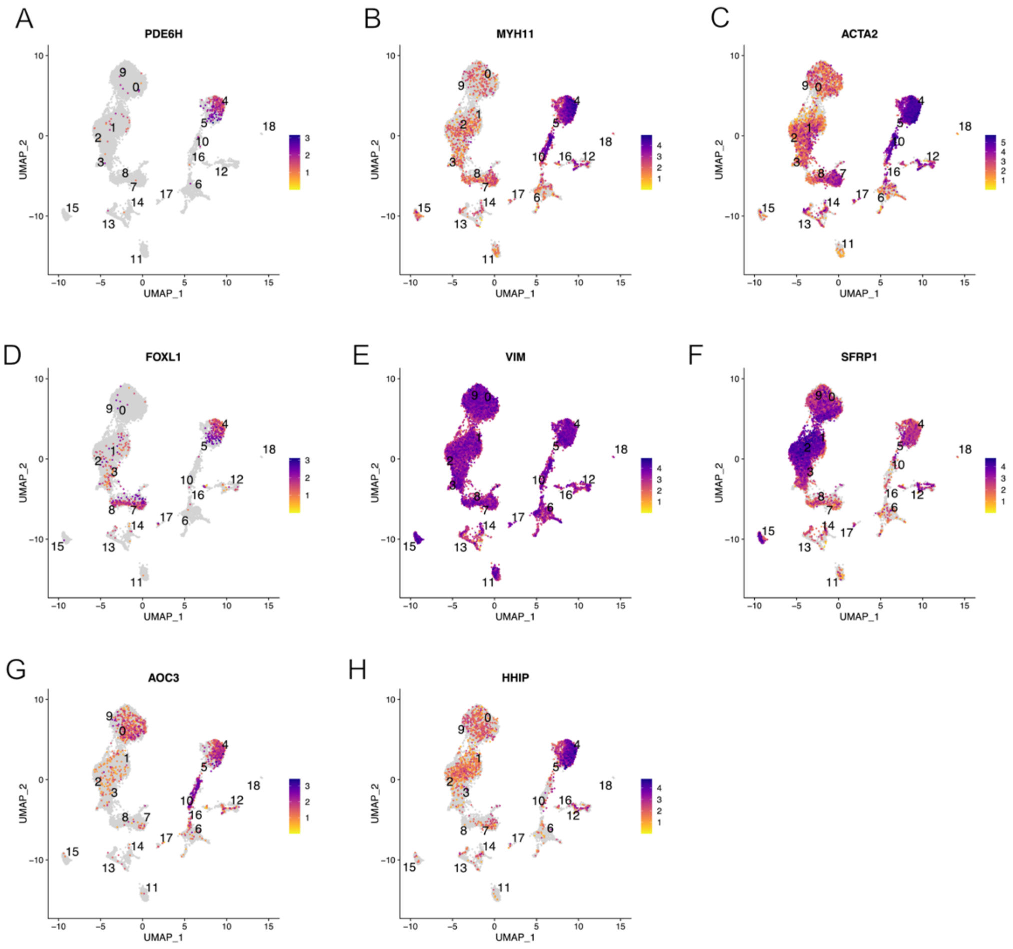


**Supplementary figure 11. Myofibroblasts**

**(A-H)** Expression of different markers of myofibroblasts projected onto UMAP plots.

### Smooth muscle/pericytes-like fibroblasts

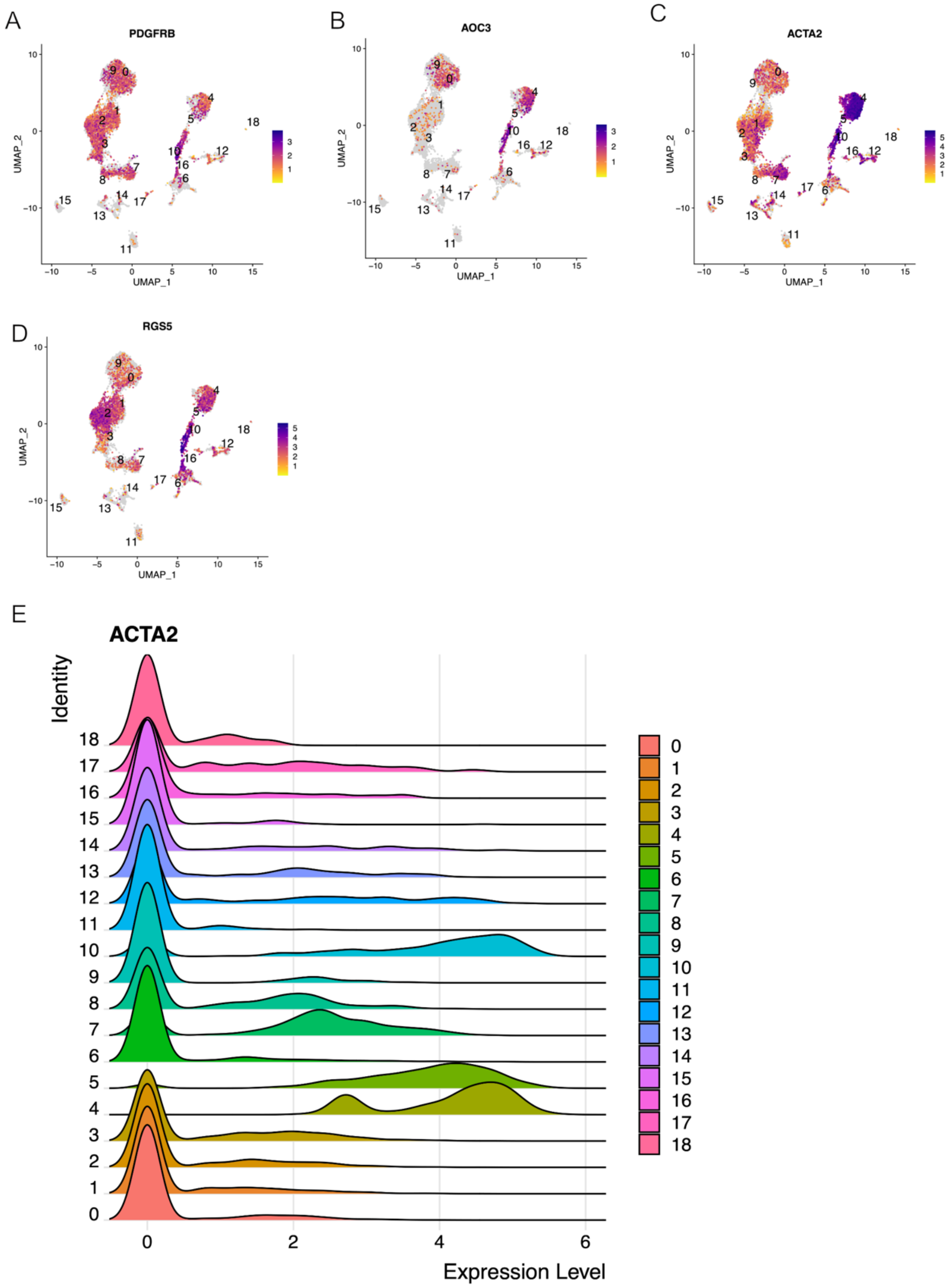


**Supplementary figure 12. Smooth muscle (SM)/ pericyte- like fibroblasts**

**(A-D)** Expression of different markers of smooth-muscle/pericyte-like fibroblasts projected onto UMAP plots.
**E)** Rigde Plot of *Acta2* expression across different fibroblast clusters.

### Supplementary Movies

**Link for the movies:**
**https://www.dropbox.com/scl/fo/jucpit1ey50qrs9rq5hrj/AFEOgI7x5qkOvAmUMvEqifY?rlkey=9uhwfhnpidc6rar9s8w05gqvb&dl=0**

**Supplementary Movie 1. Spatial organization of EGFP+ fibroblasts.**

Representation of the spindle and stellate EGFP+ fibroblasts (in green) in the distal colon sections from αSMA:CreER$^{T2}$; R26$^{mT/mG}$ mice. Other cell membranes are in red (Tomato).

**Supplementary Movie 2. EGFP+ and CD34+ fibroblasts spatial organization.**

Representation of the spindle and stellate EGFP+ fibroblasts (in green), as well as stellate CD34+ fibroblasts (in magenta) in the distal colon sections from αSMA:CreER$^{T2}$; R26$^{mT/mG}$ mice. Other cell membranes are in grey (Tomato). Stellate fibroblasts make the basket around the crypts.

**Supplementary Movie 3. EGFP+ and CD34+ fibroblasts spatial organization.**

Representation of the spindle and stellate EGFP+ fibroblasts (in green), as well as stellate CD34+ fibroblasts (in magenta) in the distal colon sections from αSMA:CreER$^{T2}$; R26$^{mT/mG}$ mice. Other cell membranes are in grey (Tomato), blood vessels are in yellow (CD31+). Stellate fibroblasts make the basket around the crypts.

### Supplementary Tables

| **Supplementary table 1- g:Profiler analysis** | |
|---|---|
| **Genes found using the code:** | |
| **cluster_markers <- FindAllMarkers(seuratObj,**<br>**logfc.threshold = 0.5**, #To test all possible genes (very slow), set this to 0<br>**only.pos = T)** #and set this to F | |
| The list of genes for each cluster was then used in the g:Profiler analysis. It can be accessed at the following links as an input for the analysis. | |
| **Reference:** | |
| Liis Kolberg, Uku Raudvere, Ivan Kuzmin, Priit Adler, Jaak Vilo, Hedi Peterson: **g:Profiler—interoperable web service for functional enrichment analysis and gene identifier mapping (2023 update)** Nucleic Acids Research, May 2023; doi:10.1093/nar/gkad347 | |
| **Results** | |
| **Cluster 0** | |
| Gprofiler link: | https://biit.cs.ut.ee/gplink/l/0ZULZSI2S6 |
| Older version: | https://biit.cs.ut.ee/gplink/l/Jo7Gw43oSO. |
| **Cluster 2** | |

| | |
|---|---|
| Gprofiler link: | https://biit.cs.ut.ee/gplink/l/cYVUyxGwTG |
| Older version: | https://biit.cs.ut.ee/gplink/l/xOdvd6PjQ8 |
| **Cluster 3** | |
| Gprofiler link: | https://biit.cs.ut.ee/gplink/l/bZyh2ym8R4 |
| Older version: | https://biit.cs.ut.ee/gplink/l/cy2P_YjxRy |
| **Cluster 4** | |
| Gprofiler link: | https://biit.cs.ut.ee/gplink/l/XOOXLXoxSq |
| Older version: | https://biit.cs.ut.ee/gplink/l/R_rhD3eKTs |
| **Cluster 5** | |
| Gprofiler link: | https://biit.cs.ut.ee/gplink/l/cdfqmDf2Rp |
| Older version: | https://biit.cs.ut.ee/gplink/l/RyZSg_E4SV |

**Supplementary Table 1.** Results of g:Profiler analysis.

#### Supplementary table 2- gene list used for creating the heatmap in Figure 1C

| p_val | avg_log2FC | pct.1 | pct.2 | p_val_adj | cluster | gene |
|---|---|---|---|---|---|---|
| **1.10429742501845E-116** | 5.65270938172795 | 0.844 | 0.032 | 3.46440188176787E-112 | 0 | Pi16 |
| **7.23244615958406E-114** | 4.06855668483596 | 0.924 | 0.08 | 2.26896300918471E-109 | 0 | Col14a1 |
| **1.97295171333568E-112** | 6.06763167678399 | 0.777 | 0.014 | 6.1895441150767E-108 | 0 | Lbp |
| **1.59280312195341E-111** | 3.89327820452841 | 0.943 | 0.114 | 4.99694195419223E-107 | 0 | Serpinf1 |
| **6.732493173638E-105** | 4.34682890441293 | 0.81 | 0.041 | 2.11211775843371E-100 | 0 | Cpxm1 |
| **9.47716552028445E-100** | 3.48739737223518 | 0.953 | 0.169 | 2.97317636702364E-95 | 0 | Cd248 |
| **1.55986160008265E-98** | 3.50498215787906 | 0.91 | 0.126 | 4.89359781177929E-94 | 0 | Plat |
| **5.48990816725031E-97** | 3.2184828243558 | 0.972 | 0.197 | 1.72229399022977E-92 | 0 | C3 |
| **3.18248951743821E-96** | 3.66134414903998 | 0.91 | 0.131 | 9.98410611410715E-92 | 0 | Htra3 |
| **2.55666635499048E-95** | 3.46119516485176 | 0.882 | 0.115 | 8.02077368887615E-91 | 0 | Lox |
| **1.99782575709296E-93** | 3.11300001822935 | 0.986 | 0.318 | 6.26757896515203E-89 | 0 | Fbn1 |
| **4.07332325393351E-93** | 3.09878582632889 | 0.962 | 0.19 | 1.27788297122402E-88 | 0 | Fbln2 |
| **2.44315506551837E-89** | 3.31203238010104 | 0.777 | 0.055 | 7.66466607154423E-85 | 0 | Dpep1 |
| **6.09517014706975E-88** | 3.23580457099875 | 0.962 | 0.327 | 1.91217677853872E-83 | 0 | Meg3 |
| **6.5398558105011E-88** | 3.17045866717502 | 0.957 | 0.279 | 2.0516835648704E-83 | 0 | Mfap5 |
| **4.01124724325598E-87** | 2.83410784189517 | 0.991 | 0.307 | 1.25840848515427E-82 | 0 | Igfbp4 |
| **1.08806781909347E-86** | 2.95121187327177 | 0.948 | 0.202 | 3.41348636206003E-82 | 0 | Igfbp6 |
| **4.17578009674414E-85** | 3.25346142911363 | 0.877 | 0.151 | 1.31002573195057E-80 | 0 | Prss23 |
| **1.69154190353278E-83** | 2.82208080176691 | 0.891 | 0.139 | 5.30670525976303E-79 | 0 | Cd34 |
| **5.05555542776185E-83** | 2.5534990352948 | 1 | 0.565 | 1.58602884879745E-78 | 0 | Fstl1 |
| **1.22155883217233E-82** | 2.61553864304553 | 0.981 | 0.371 | 3.83227436829104E-78 | 0 | Ccdc80 |
| **1.48843194074683E-77** | 2.83538189455714 | 0.844 | 0.14 | 4.66950868451096E-73 | 0 | Man2a1 |
| **1.52377551554345E-77** | 2.35109651814587 | 0.995 | 0.837 | 4.78038854736292E-73 | 0 | Col3a1 |
| **1.72421706599293E-77** | 2.48249465319025 | 0.953 | 0.277 | 5.40921377943303E-73 | 0 | Aebp1 |
| **7.69012300869433E-75** | 2.51514723184996 | 0.962 | 0.329 | 2.41254539028759E-70 | 0 | Nid1 |
| **9.77825804194993E-75** | 1.89889347249068 | 1 | 0.865 | 3.06763511292053E-70 | 0 | Sparc |
| **2.7089361988663E-74** | 2.46701139652503 | 0.91 | 0.195 | 8.49847464308336E-70 | 0 | Ly6c1 |
| **4.14391830199869E-74** | 2.3291690588252 | 0.976 | 0.298 | 1.30003004970303E-69 | 0 | Tnxb |
| **9.70967247720645E-74** | 3.45544398780389 | 0.867 | 0.206 | 3.04611844954921E-69 | 0 | Mt2 |
| **8.08781552850887E-72** | 3.20692686983762 | 0.891 | 0.22 | 2.5373094876038E-67 | 0 | Eln |
| **2.84940534266338E-71** | 2.35507992940342 | 0.919 | 0.249 | 8.93915444100356E-67 | 0 | Loxl1 |
| **4.96790777103259E-71** | 2.38544036928523 | 0.81 | 0.117 | 1.55853202592834E-66 | 0 | Sulf2 |
| **1.27530802792682E-69** | 2.54142778931578 | 0.806 | 0.119 | 4.00089634521203E-65 | 0 | Angptl4 |
| **2.88790243010598E-68** | 3.06094402667701 | 0.773 | 0.124 | 9.05992750372847E-64 | 0 | Matn2 |
| **6.21636301342329E-67** | 2.2962978232085 | 0.943 | 0.295 | 1.95019740457115E-62 | 0 | Clec3b |
| **9.56085507256133E-67** | 2.81531584306195 | 0.844 | 0.185 | 2.99943145336394E-62 | 0 | Gas1 |
| **7.85528511570717E-66** | 2.34370925025955 | 0.844 | 0.162 | 2.46436004649965E-61 | 0 | Heg1 |

| | | | | | | |
|---|---|---|---|---|---|---|
| **1.19592648719004E-65** | 2.584434321115 | 0.787 | 0.133 | 3.75186057561261E-61 | 0 | Slit3 |
| **1.23077494332173E-65** | 2.48212829124891 | 0.877 | 0.24 | 3.86118715218893E-61 | 0 | Lsp1 |
| **2.47834524637043E-63** | 1.94979301231103 | 0.972 | 0.394 | 7.77506470691332E-59 | 0 | Dpt |
| **3.82801188969414E-63** | 2.22497347410783 | 0.877 | 0.213 | 1.20092389003484E-58 | 0 | Mmp23 |
| **2.24527573869946E-62** | 2.49571910853392 | 0.768 | 0.144 | 7.04387904744793E-58 | 0 | Ctsh |
| **3.71852445460218E-62** | 1.79714218308367 | 0.986 | 0.65 | 1.1665754918978E-57 | 0 | Dcn |
| **3.97182016689867E-62** | 2.05793901518797 | 0.967 | 0.369 | 1.24603942275945E-57 | 0 | Lrp1 |
| **1.11760219714983E-61** | 2.05176207431939 | 0.957 | 0.449 | 3.50614161289845E-57 | 0 | Ly6a |
| **1.21055964787254E-61** | 2.44670523275171 | 0.806 | 0.162 | 3.79776772730575E-57 | 0 | Timp1 |
| **1.89147552774601E-61** | 2.73213125335579 | 0.754 | 0.128 | 5.93393702564477E-57 | 0 | Cd55 |
| **1.04893352959154E-60** | 2.00888981451739 | 0.972 | 0.414 | 3.29071426903459E-56 | 0 | Lum |
| **1.14135828355519E-60** | 2.13909625369169 | 0.844 | 0.179 | 3.58066920716936E-56 | 0 | Islr |
| **3.27261753485694E-59** | 1.82340672361992 | 0.986 | 0.622 | 1.02668557303532E-54 | 0 | Serping1 |
| **1.49349199919341E-58** | 1.93585702257157 | 0.995 | 0.694 | 4.68538309986956E-54 | 0 | Col1a1 |
| **1.57847483139576E-56** | 1.94722813043424 | 0.863 | 0.202 | 4.95199124105478E-52 | 0 | Bmp1 |
| **1.91840876125724E-56** | 2.27875313593502 | 0.787 | 0.171 | 6.01843196581621E-52 | 0 | Mfap2 |
| **6.88407900566685E-56** | 1.85413896951251 | 0.758 | 0.128 | 2.15967326565781E-51 | 0 | C4b |
| **2.55400362987914E-54** | 2.31354020895495 | 0.777 | 0.176 | 8.01242018765682E-50 | 0 | Adamts2 |
| **2.89317195253194E-54** | 1.97567864448993 | 0.91 | 0.346 | 9.0764590494832E-50 | 0 | Rnase4 |
| **2.56544853847156E-51** | 1.80233378860758 | 0.981 | 0.499 | 8.04832515489298E-47 | 0 | Col5a2 |
| **4.47784065333304E-51** | 1.47434517619885 | 0.915 | 0.266 | 1.40478816976364E-46 | 0 | Fbln1 |
| **4.88025705516673E-51** | 1.8533500073403 | 0.82 | 0.21 | 1.53103424334691E-46 | 0 | BC042477 |
| **5.69558256664272E-51** | 1.78018817460882 | 0.773 | 0.16 | 1.78681816280715E-46 | 0 | Thbs3 |
| **1.57494388551312E-50** | 1.67273939592307 | 0.858 | 0.229 | 4.94091395763175E-46 | 0 | Vcan |
| **1.863434029602E-50** | 1.48874524872249 | 0.991 | 0.65 | 5.84596523766741E-46 | 0 | Gsn |
| **3.86895792364615E-50** | 1.97126189426947 | 0.796 | 0.215 | 1.21376947980627E-45 | 0 | Fxyd5 |
| **3.82875415433977E-45** | 1.64641754213091 | 0.915 | 0.366 | 1.20115675329947E-40 | 0 | C1ra |
| **9.02134277122461E-45** | 1.57389561423989 | 0.953 | 0.391 | 2.83017565418858E-40 | 0 | Col5a1 |
| **2.38240953233929E-44** | 1.49822274765239 | 0.962 | 0.574 | 7.47409518485483E-40 | 0 | Ctsl |
| **6.55700259283741E-43** | 1.88718564786828 | 0.915 | 0.377 | 2.05706285342495E-38 | 0 | Plpp3 |
| **8.83564201006365E-42** | 1.27987084040552 | 0.863 | 0.249 | 2.77191761139717E-37 | 0 | Tgfbr3 |
| **1.726917856236E-41** | 1.48497001751105 | 0.905 | 0.334 | 5.41768669858358E-37 | 0 | Tgfbr2 |
| **2.06371472271799E-41** | 1.65259735562136 | 0.777 | 0.25 | 6.47428582811088E-37 | 0 | S100a16 |
| **3.57285642482252E-41** | 1.32134202988623 | 0.995 | 0.803 | 1.12087651759532E-36 | 0 | Col1a2 |
| **3.65784172431083E-40** | 1.29326917650294 | 0.801 | 0.224 | 1.14753810575079E-35 | 0 | Olfml2b |
| **1.94863291144983E-39** | 1.28408290306588 | 0.905 | 0.306 | 6.11325116980041E-35 | 0 | Cd81 |
| **3.72334676834837E-39** | 1.17278211950542 | 0.957 | 0.414 | 1.16808834816625E-34 | 0 | Mmp14 |
| **2.95972782371446E-38** | 1.48165893237755 | 0.886 | 0.337 | 9.285258128557E-34 | 0 | P4hb |
| **3.48985217805949E-38** | 1.88529844685578 | 0.853 | 0.343 | 1.09483642530082E-33 | 0 | Mt1 |
| **1.50354957336213E-37** | 1.3309523557079 | 0.943 | 0.567 | 4.71693572155167E-33 | 0 | Timp2 |
| **3.33572924487089E-37** | 1.31553885136442 | 0.905 | 0.348 | 1.04648497870089E-32 | 0 | Mmp2 |
| **1.52003561865758E-36** | 1.48021993648796 | 0.754 | 0.224 | 4.76865574285255E-32 | 0 | Pbx1 |
| **4.31356496567651E-36** | 1.2473249515631 | 0.905 | 0.382 | 1.35325160103204E-31 | 0 | Pdia6 |
| **4.71698874634297E-36** | 1.45603977483696 | 0.787 | 0.245 | 1.47981370950272E-31 | 0 | Lamb2 |
| **6.70002178813137E-36** | 1.53926962205205 | 0.754 | 0.243 | 2.10193083537257E-31 | 0 | Tmed3 |
| **1.30809499487795E-33** | 1.16590910926983 | 0.872 | 0.323 | 4.10375561793111E-29 | 0 | Spon2 |
| **6.69817389938927E-32** | 1.13056390722181 | 0.915 | 0.419 | 2.1013511157164E-27 | 0 | Lamc1 |
| **2.230080539889E-31** | 1.09771584425807 | 0.957 | 0.581 | 6.99620866973977E-27 | 0 | Calr |
| **4.47077443845019E-31** | 1.01394520393684 | 0.815 | 0.279 | 1.40257135683059E-26 | 0 | Txndc5 |
| **3.83131259005712E-29** | 1.21550191322106 | 0.758 | 0.286 | 1.20195938575272E-24 | 0 | Cnpy2 |
| **2.47448732311321E-28** | 1.23869591870652 | 0.839 | 0.369 | 7.76296163007077E-24 | 0 | Axl |
| **4.20020053035381E-25** | 1.13433302986887 | 0.834 | 0.378 | 1.3176869103826E-20 | 0 | Serpinb6a |
| **4.77502659669702E-23** | 1.16294460560146 | 0.896 | 0.535 | 1.49802134391579E-18 | 0 | Pcolce |
| **9.47238549003783E-23** | 1.00745722173549 | 0.806 | 0.362 | 2.97167677593467E-18 | 0 | S100a13 |
| **7.8782912082042E-67** | 4.09378181364869 | 0.82 | 0.169 | 2.47157751783782E-62 | 2 | Procr |
| **7.54089886970791E-62** | 2.77214686057625 | 0.773 | 0.115 | 2.36573079340476E-57 | 2 | Bmp5 |
| **1.77546076727406E-56** | 2.74327254099652 | 0.93 | 0.339 | 5.5699755190922E-52 | 2 | Cxcl12 |
| **1.21321848056112E-47** | 2.5289924922689 | 0.812 | 0.22 | 3.80610901721633E-43 | 2 | Bmp4 |
| **7.6945028662691E-47** | 2.7625163458973 | 0.844 | 0.313 | 2.41391943920594E-42 | 2 | Tcf4 |
| **3.65477591655567E-46** | 2.43142698569393 | 0.906 | 0.375 | 1.14657630054184E-41 | 2 | Pdgfra |
| **1.08110881443472E-44** | 2.6100513108655 | 0.789 | 0.217 | 3.39165457264461E-40 | 2 | Ifitm1 |
| **1.09230718073933E-43** | 2.55318985884461 | 0.844 | 0.319 | 3.42678608741543E-39 | 2 | Tmem119 |
| **3.13020959790013E-41** | 2.27111351392798 | 0.797 | 0.22 | 9.82009355053228E-37 | 2 | Adamdec1 |
| **4.89189479068517E-35** | 1.50932789178021 | 0.969 | 0.574 | 1.53468523373375E-30 | 2 | Gpx3 |
| **5.8994349291679E-34** | 1.46783274991505 | 0.969 | 0.663 | 1.85077072597855E-29 | 2 | Col6a1 |
| **4.39183945466567E-31** | 1.21946904707911 | 0.992 | 0.876 | 1.37780787371771E-26 | 2 | Tmsb10 |
| **9.99916813644152E-31** | 1.92570547707412 | 0.852 | 0.517 | 3.13693902776443E-26 | 2 | Nbl1 |
| **1.13180000353169E-29** | 1.44153188111311 | 0.953 | 0.676 | 3.55068297107963E-25 | 2 | Col6a2 |
| **1.10334875267766E-26** | 1.62433461324224 | 0.883 | 0.548 | 3.46142570690036E-22 | 2 | Ogn |
| **7.44060802541254E-24** | 1.69246566929245 | 0.797 | 0.477 | 2.33426754973242E-19 | 2 | Lamb1 |
| **1.2089491272029E-23** | 1.64235835919226 | 0.812 | 0.481 | 3.79271520186094E-19 | 2 | Htra1 |

| | | | | | | |
|---|---|---|---|---|---|---|
| **1.7459126980272E-22** | 1.08508011506274 | 0.953 | 0.788 | 5.47727731625093E-18 | 2 | Cst3 |
| **1.30778846612197E-20** | 1.56115309650607 | 0.789 | 0.458 | 4.10279397591785E-16 | 2 | Pmepa1 |
| **2.58665476429381E-19** | 1.23339969694409 | 0.828 | 0.435 | 8.11485332654253E-15 | 2 | Mmp2 |
| **3.07393834430069E-14** | 1.01560868331474 | 0.867 | 0.689 | 9.64355937374012E-10 | 2 | Ifitm2 |
| **4.80316876995777E-13** | 1.22753105394318 | 0.766 | 0.526 | 1.50685010651115E-08 | 2 | Col6a3 |
| **5.78308867810868E-13** | 1.18275570947245 | 0.758 | 0.579 | 1.81427058009625E-08 | 2 | Myl12a |
| **2.64082613048691E-121** | 7.27769795804054 | 0.893 | 0.041 | 8.28479973656353E-117 | 3 | Hhip |
| **4.24399919960821E-58** | 2.67912303369707 | 0.901 | 0.208 | 1.33142742890109E-53 | 3 | Myh11 |
| **1.59463008974151E-55** | 3.69897032344404 | 0.785 | 0.173 | 5.00267351753707E-51 | 3 | F2r |
| **7.57451798107452E-39** | 2.34654567129557 | 0.917 | 0.582 | 2.3762777810227E-34 | 3 | Flna |
| **7.44964057997531E-38** | 1.84512656171704 | 0.901 | 0.361 | 2.33710124274985E-33 | 3 | Myl9 |
| **4.19052464267465E-37** | 2.61213668965535 | 0.826 | 0.387 | 1.31465139089989E-32 | 3 | Mylk |
| **1.12486942343626E-35** | 1.6717725784662 | 0.942 | 0.441 | 3.52894035520424E-31 | 3 | Acta2 |
| **2.88055353963969E-34** | 1.60104843171711 | 0.893 | 0.332 | 9.03687256455765E-30 | 3 | Tagln |
| **5.54022286023882E-33** | 1.87241895594595 | 0.917 | 0.576 | 1.73807871571412E-28 | 3 | Tmem176b |
| **5.94267604819871E-32** | 2.24065451645445 | 0.76 | 0.302 | 1.8643363298409E-27 | 3 | Ckb |
| **7.67392996765375E-32** | 1.74732537210438 | 0.926 | 0.556 | 2.40746530945233E-27 | 3 | Tpm1 |
| **3.04797928444491E-30** | 1.78976521913037 | 0.876 | 0.507 | 9.56212061116057E-26 | 3 | Tpm2 |
| **6.56721452015699E-27** | 1.92401241558264 | 0.818 | 0.478 | 2.06026653926365E-22 | 3 | Tmem176a |
| **8.06180944678504E-26** | 1.36118132531153 | 0.909 | 0.801 | 2.5291508596454E-21 | 3 | Myl6 |
| **1.39510098302371E-25** | 1.76109015159807 | 0.851 | 0.658 | 4.376710803942E-21 | 3 | Dstn |
| **1.0771121459919E-23** | 1.41478663594285 | 0.909 | 0.596 | 3.37911622440577E-19 | 3 | Ltbp4 |
| **5.39126034890877E-22** | 1.12728725635238 | 0.95 | 0.628 | 1.69134619665966E-17 | 3 | Postn |
| **1.31647207207975E-16** | 1.24244541180844 | 0.802 | 0.387 | 4.13003618452858E-12 | 3 | EGFP |
| **8.00793994321883E-14** | 1.27731111335907 | 0.785 | 0.678 | 2.51225091898661E-09 | 3 | Itgb1 |
| **2.30747052179139E-13** | 1.03770094867726 | 0.851 | 0.744 | 7.23899652096394E-09 | 3 | Rarres2 |
| **1.61630333695036E-12** | 1.05444490542422 | 0.785 | 0.623 | 5.07066682868068E-08 | 3 | Plac8 |
| **4.28009852543672E-11** | 1.09921103994454 | 0.752 | 0.6 | 1.34275250940001E-06 | 3 | Prkcdbp |
| **3.47104019994855E-122** | 7.78180821263112 | 0.783 | 0.006 | 1.08893473152786E-117 | 4 | Notch3 |
| **1.65792911780942E-110** | 6.48790306634289 | 0.783 | 0.019 | 5.20125522839172E-106 | 4 | Cox4i2 |
| **3.8646618560931E-105** | 5.84214896359246 | 0.84 | 0.043 | 1.21242171749353E-100 | 4 | Ndufa4l2 |
| **1.13326649704191E-100** | 4.89368168664934 | 0.83 | 0.042 | 3.55528365451986E-96 | 4 | Bcam |
| **1.73649620815841E-100** | 5.0454849793582 | 0.887 | 0.064 | 5.44773590423455E-96 | 4 | Rgs5 |
| **2.47953853059602E-98** | 4.55204438520172 | 0.906 | 0.076 | 7.77880827818582E-94 | 4 | Mef2c |
| **2.87717616722441E-78** | 4.09397528425379 | 0.858 | 0.114 | 9.02627707181643E-74 | 4 | Des |
| **1.27735335792878E-62** | 3.1493173805021 | 0.792 | 0.105 | 4.00731295449418E-58 | 4 | Tinagl1 |
| **2.56212794710802E-58** | 3.12486894872035 | 0.783 | 0.115 | 8.03790779566729E-54 | 4 | Epas1 |
| **1.12509147471991E-55** | 2.82167069934496 | 0.972 | 0.332 | 3.52963697449129E-51 | 4 | Tagln |
| **6.6153164801163E-53** | 3.16587979222634 | 0.925 | 0.316 | 2.07535708614209E-48 | 4 | Mustn1 |
| **4.67954376095055E-51** | 3.02068792866443 | 0.915 | 0.352 | 1.46806646868541E-46 | 4 | Tm4sf1 |
| **1.66093143170225E-50** | 2.68024042143011 | 0.981 | 0.446 | 5.2106740875363E-46 | 4 | Acta2 |
| **3.93336503953271E-47** | 2.04595170173219 | 0.925 | 0.22 | 1.2339752802022E-42 | 4 | Myh11 |
| **2.88746959375622E-45** | 2.60791857196447 | 0.877 | 0.287 | 9.05856960953202E-41 | 4 | Ppp1r12a |
| **3.95349342323441E-45** | 2.51153084524424 | 0.943 | 0.367 | 1.2402899567371E-40 | 4 | Myl9 |
| **1.35975790940873E-43** | 2.22146895097757 | 0.972 | 0.618 | 4.26583251339706E-39 | 4 | Sparcl1 |
| **3.84502350805723E-42** | 2.23110848339525 | 0.962 | 0.648 | 1.20626077494771E-37 | 4 | Cald1 |
| **9.63475588291007E-41** | 2.45567522818187 | 0.802 | 0.222 | 3.02261561558655E-36 | 4 | Ccnd2 |
| **4.65212617339711E-40** | 2.9102354739343 | 0.783 | 0.244 | 1.45946502311814E-35 | 4 | Gng11 |
| **4.41996320631861E-39** | 2.2495424446542 | 0.953 | 0.503 | 1.38663085708627E-34 | 4 | Tpm2 |
| **2.6187127979638E-33** | 2.08567585983331 | 0.83 | 0.344 | 8.21542578977203E-29 | 4 | Ppp1cb |
| **3.41596265345818E-33** | 2.56188233627096 | 0.774 | 0.278 | 1.0716558036429E-28 | 4 | Oaz2 |
| **4.29935272973875E-33** | 2.03989084880353 | 0.821 | 0.293 | 1.34879293837364E-28 | 4 | Mgst3 |
| **5.41115583725294E-33** | 1.78921485043035 | 0.943 | 0.561 | 1.69758780926299E-28 | 4 | Tpm1 |
| **6.26131255789758E-33** | 1.88867611916362 | 0.981 | 0.9 | 1.96429897566363E-28 | 4 | Crip1 |
| **1.72690082147106E-32** | 1.98063388429986 | 0.896 | 0.47 | 5.417633257119E-28 | 4 | Mfge8 |
| **3.04004388130662E-32** | 1.67523011371215 | 0.943 | 0.606 | 9.53722566443512E-28 | 4 | Calm2 |
| **3.37705469906718E-32** | 1.84082566170274 | 0.915 | 0.379 | 1.05944960019136E-27 | 4 | EGFP |
| **1.07523244439054E-31** | 1.64066161540124 | 0.972 | 0.793 | 3.37321922454202E-27 | 4 | Myl6 |
| **1.09183011536237E-30** | 1.78149083419773 | 0.887 | 0.413 | 3.42528943791481E-26 | 4 | Crip2 |
| **2.05003193041977E-29** | 2.07775281585972 | 0.802 | 0.305 | 6.43136017211291E-25 | 4 | Ckb |
| **1.98623017945736E-27** | 1.87971823234661 | 0.811 | 0.361 | 6.23120131899364E-23 | 4 | Prkar1a |
| **2.07316560530462E-24** | 1.43127091628125 | 0.906 | 0.612 | 6.50393513696165E-20 | 4 | Calm1 |
| **6.84514956932344E-24** | 1.71170972617584 | 0.83 | 0.377 | 2.14746032288815E-19 | 4 | Filip1l |
| **1.58475122539176E-22** | 1.09169610612434 | 0.887 | 0.388 | 4.97168154429904E-18 | 4 | Mylk |
| **2.24345907119049E-22** | 1.12248083007053 | 0.962 | 0.722 | 7.0381797981388E-18 | 4 | Cox8a |
| **3.37369748889234E-22** | 1.62204567204972 | 0.83 | 0.518 | 1.05839637621531E-17 | 4 | Fxyd1 |
| **1.37733614838881E-21** | 1.58451691125882 | 0.755 | 0.322 | 4.32097896472536E-17 | 4 | Actn1 |
| **7.01141309085255E-21** | 1.7472852216885 | 0.755 | 0.377 | 2.19962051486226E-16 | 4 | GSMG0070069 |
| **1.49794430196159E-19** | 1.34131502201708 | 0.887 | 0.657 | 4.6993508641139E-15 | 4 | Dstn |
| **3.55066093624276E-19** | 1.05738953712244 | 1 | 1 | 1.11391334891808E-14 | 4 | Vim |
| **1.20256734382743E-18** | 1.50432310691948 | 0.802 | 0.452 | 3.77269427105542E-14 | 4 | sept-07 |

| | | | | | | |
|---|---|---|---|---|---|---|
| **2.47204047151482E-18** | 1.11916429855927 | 0.811 | 0.371 | 7.7552853672363E-14 | 4 | Timp3 |
| **5.77298910518415E-18** | 1.43342386181624 | 0.802 | 0.548 | 1.81110214207837E-13 | 4 | Slc25a4 |
| **1.75258968542594E-17** | 1.2679885305263 | 0.849 | 0.609 | 5.49822436111826E-13 | 4 | Atp5b |
| **4.78814455005956E-16** | 1.15102792271543 | 0.849 | 0.629 | 1.50213670824469E-11 | 4 | Ubb |
| **7.2765020177532E-16** | 1.03933186926437 | 0.849 | 0.645 | 2.28278421300953E-11 | 4 | Chchd2 |
| **6.42541098001804E-15** | 1.18521218239993 | 0.792 | 0.458 | 2.01577993265126E-10 | 4 | Tsc22d1 |
| **1.56986766662413E-14** | 1.47433690933061 | 0.792 | 0.457 | 4.92498884373321E-10 | 4 | Id3 |
| **1.98613227388744E-14** | 1.12204789251757 | 0.858 | 0.705 | 6.23089416963969E-10 | 4 | Dynll1 |
| **1.98811445254938E-14** | 1.08060861055029 | 0.764 | 0.398 | 6.2371126605379E-10 | 4 | Eif4a2 |
| **2.67680985002846E-14** | 1.10600675294358 | 0.877 | 0.581 | 8.39768786150929E-10 | 4 | Atpif1 |
| **4.32546149846573E-14** | 1.03440973786589 | 0.783 | 0.488 | 1.35698378129867E-09 | 4 | Slc25a5 |
| **1.54487589277272E-13** | 1.01829018958726 | 0.83 | 0.591 | 4.84658465080657E-09 | 4 | Prkcdbp |
| **2.02673840695781E-13** | 1.03819372405162 | 0.991 | 0.919 | 6.35828373030804E-09 | 4 | GSMG0065070 |
| **3.16662930633813E-13** | 1.15367432297819 | 0.774 | 0.448 | 9.93434945984399E-09 | 4 | Fus |
| **1.9043605231266E-12** | 1.11425763094724 | 0.934 | 0.757 | 5.97435983315278E-08 | 4 | GSMG0067152 |
| **1.91002912378668E-10** | 1.12735330628305 | 0.764 | 0.49 | 5.99214336714358E-06 | 4 | Tppp3 |
| **4.30854654625092E-103** | 6.52996070375483 | 0.784 | 0.011 | 1.35167722248984E-98 | 5 | Myrf |
| **6.12749981053074E-103** | 8.68802506936619 | 0.784 | 0.011 | 1.9223192405597E-98 | 5 | Slpi |
| **1.80818283061083E-62** | 5.18051780192148 | 0.757 | 0.038 | 5.67263117619231E-58 | 5 | Cxadr |
| **2.93290635499962E-50** | 5.13867160806492 | 0.865 | 0.088 | 9.20111381690481E-46 | 5 | Clu |
| **5.11453609596312E-31** | 3.68876733381158 | 0.919 | 0.206 | 1.60453226402555E-26 | 5 | AK088706 |
| **1.8248577357711E-25** | 3.23920333963443 | 0.919 | 0.258 | 5.72494368866109E-21 | 5 | Ptgis |
| **1.4382763893209E-24** | 2.83922569580251 | 0.919 | 0.252 | 4.51216068857753E-20 | 5 | LF200403 |
| **2.19787189218064E-21** | 3.06011486240313 | 0.757 | 0.176 | 6.8951637001491E-17 | 5 | AK075710 |
| **5.09168538268559E-19** | 2.10192998060683 | 0.973 | 0.346 | 1.59736353825612E-14 | 5 | BC042477 |
| **3.58802341263399E-17** | 2.54699129510538 | 0.811 | 0.242 | 1.12563470501154E-12 | 5 | Cfb |
| **4.31008744843517E-15** | 1.60257029031476 | 1 | 0.379 | 1.35216063432308E-10 | 5 | C3 |
| **1.36263182543804E-14** | 1.74454456239373 | 1 | 0.963 | 4.27484856276423E-10 | 5 | Fth1 |
| **1.13878868563421E-12** | 2.28281715863376 | 0.784 | 0.303 | 3.57260786457165E-08 | 5 | Sdc4 |
| **2.28612799386103E-12** | 1.96786227687599 | 0.892 | 0.357 | 7.17204074234081E-08 | 5 | Csrp2 |
| **2.15341067124234E-11** | 1.60570109569517 | 0.838 | 0.323 | 6.75567995782147E-07 | 5 | Heg1 |
| **1.16200788237116E-09** | 1.31576943490997 | 0.757 | 0.27 | 3.64545112857479E-05 | 5 | Crtap |
| **6.26196964236272E-09** | 1.08444460211527 | 1 | 0.782 | 0.000196450511620203 | 5 | B2m |
| **2.05063886311795E-08** | 1.10342679475447 | 0.865 | 0.468 | 0.000643326424137363 | 5 | P4hb |
| **5.05660599178988E-08** | 1.57344018712063 | 0.892 | 0.65 | 0.00158635843174432 | 5 | H2-K1 |
| **8.53193588005821E-08** | 1.29601766269719 | 0.757 | 0.319 | 0.00267663892429186 | 5 | Mlec |
| **9.39527503759086E-08** | 1.25025392077389 | 0.838 | 0.499 | 0.002947485684793 | 5 | Npc2 |
| **1.20130191710438E-07** | 1.14159054645276 | 0.892 | 0.44 | 0.00376872437433985 | 5 | Aebp1 |
| **1.93804517535205E-07** | 1.33418061482106 | 0.838 | 0.494 | 0.00608003532411444 | 5 | Pabpc1 |
| **2.00912355829263E-07** | 1.07100523091634 | 0.784 | 0.361 | 0.00630302242707565 | 5 | Atp1a1 |
| **2.53253854197421E-07** | 1.07756114255363 | 0.892 | 0.448 | 0.00794507991388148 | 5 | Cd81 |
| **5.55034841496515E-07** | 1.2484731946511 | 0.757 | 0.403 | 0.0174125530474287 | 5 | Ssr3 |
| **1.55829578180844E-06** | 1.2265010928345 | 0.784 | 0.414 | 0.0488868552668944 | 5 | Timp3 |

**Supplementary Table 2.** List of the markers for each fibroblast population (logFC=1, genes expressed in at least 75% of cells, with p_val_adj <= 0.05) used for creating the heatmap in Figure 1C.
